# Language models generalize beyond natural proteins

**DOI:** 10.1101/2022.12.21.521521

**Authors:** Robert Verkuil, Ori Kabeli, Yilun Du, Basile I. M. Wicky, Lukas F. Milles, Justas Dauparas, David Baker, Sergey Ovchinnikov, Tom Sercu, Alexander Rives

**Affiliations:** Meta Fundamental AI Research Protein Team (FAIR); Massachusetts Institute of Technology. Work performed as visiting researcher at Meta FAIR; Department of Biochemistry, University of Washington, Seattle, WA, USA; Institute for Protein Design, University of Washington, Seattle, WA, USA; Howard Hughes Medical Institute, University of Washington, Seattle, WA, USA; John Harvard Distinguished Science Fellowship Program, Harvard University, Cambridge, MA, USA; New York University

## Abstract

Learning the design patterns of proteins from sequences across evolution may have promise toward generative protein design. However it is unknown whether language models, trained on sequences of natural proteins, will be capable of more than memorization of existing protein families. Here we show that language models generalize beyond natural proteins to generate *de novo* proteins. We focus on two protein design tasks: fixed backbone design where the structure is specified, and unconstrained generation where the structure is sampled from the model. Remarkably although the models are trained only on sequences, we find that they are capable of designing structure. A total of 228 generated proteins are evaluated experimentally with high overall success rates (152/228 or 67%) in producing a soluble and monomeric species by size exclusion chromatography. Out of 152 experimentally successful designs, 35 have no significant sequence match to known natural proteins. Of the remaining 117, sequence identity to the nearest sequence match is at median 27%, below 20% for 6 designs, and as low as 18% for 3 designs. For fixed backbone design, the language model generates successful designs for each of eight experimentally evaluated artificially created fixed backbone targets. For unconstrained generation, sampled proteins cover diverse topologies and secondary structure compositions, and have high experimental success rate (71/129 or 55%). The designs reflect deep patterns linking sequence and structure, including motifs that occur in related natural structures, and motifs that are not observed in similar structural contexts in known protein families. The results show that language models, though only trained on sequences, learn a deep grammar that enables the design of protein structure, extending beyond natural proteins.

## Introduction

Generative artificial intelligence for biology has potential to open up a space of protein design beyond natural proteins. Since amino acid sequences are the fundamental codes of proteins, learning to read and write these codes with a language model may have promise. Language models have played a central role in recent advances in artificial intelligence (1), including developments in complex reasoning, mathematical problem solving, image generation, and natural language generation (2–4). Scaling laws link performance with the compute, data, and number of parameters used to train the models (5), and emergence of higher level capabilities is observed with increasing scale (6). In biology, recent work on evolutionary scale language models of proteins has shown that a deep knowledge of intrinsic biological properties emerges from training on protein sequences (7). Information about the folded three dimensional structure of proteins develops within the models, extending to atomic resolution structure (8). This information emerges through training on sequences alone. At the same time the structural information that emerges as a result of training on sequences has been shown to depend on the available evolutionary information, varying as a function of the number of related proteins in the training data (8, 9). It is an open question across domains to what extent language models are capable of generalizing outside their training data. In biology, it is unknown whether language models can be used to explore a design space beyond that of natural proteins.

Here we demonstrate that language models generalize beyond natural proteins to generate *de novo* proteins, different in sequence and structure from natural proteins. We experimentally validate a large number of designs spanning diverse topologies and sequences. We find that although language models are trained only on the sequences of proteins, they are capable of designing protein structure, including structures of artificially engineered de novo proteins that are distinct from those of natural proteins. Given the backbone of a de novo protein structure as a target, the language model generates sequences that are predicted to fold to the specified structure. When the sequence and structure are both free, language models produce designs that span a wide range of fold topologies and secondary structure compositions, creating proteins which overlap the natural sequence distribution as well as extend beyond it. Designs succeed experimentally across the space of sampled proteins, including many designs that are distant in sequence from natural proteins. The model generates motifs that link sequence to the design of structure and can apply them in new sequence and structural contexts, including motifs such as complex hydrogen bond networks that are not found in sequence- or structurally-similar known proteins. Overall experimental success rates are high with 152 out of a total of 228 (67%) experimentally evaluated proteins producing a soluble and monomeric species by size exclusion chromatography (SEC). The high success rate extends to proteins that are distant from natural proteins where 31 out of a total of 49 (63%) experimentally evaluated proteins succeed.

### A deep grammar of protein sequences

We hypothesize that there exists a deep underlying grammar in protein sequences that makes it possible for the language model to generalize. To generalize beyond natural proteins, language models will need to access design patterns that extend outside the space of natural proteins. Classically this form generalization has been enabled by an energy function grounded in physics that captures the native folded state (10). Recently deep learning based methods grounded in structure have been proposed as a new approach to this problem by inverting structure prediction (11, 12), or conditioning on backbone structures (13–15). By modeling the structure explicitly during training, new deep learning approaches may capture something similar to the physical energy (16). The success of language models on this problem suggests that deep patterns in sequences may offer an alternative path to generalization, independent of an explicit model of the underlying physics.

The classical perspective of evolutionary inference from sequences is that information about the properties of proteins is encoded into the sequence patterns of evolutionarily related proteins through conservation and coevolution. This view develops from the observation that the statistics of protein families reflect the constraints acting on the evolution of the sequences including biological structure and function (17, 18). This insight has formed the basis for the inference of structure and function from sequences in a protein family (19), and has also recently been applied with success by generative models to generate new examples from existing protein families (20–22). To date experimental validation of sequence based models for protein design has been limited to natural protein families.

Accessing a *de novo* design space distant from naturally occurring protein families is a fundamentally more challenging problem. This problem by definition cannot be solved by generating new samples from naturally occurring protein families. To solve this problem with a model grounded in sequences, it will be necessary to learn sequence patterns that generalize beyond individual protein families. Evolutionary scale language models go beyond classic protein family models by training on diverse sequences across evolution which means that they have the potential to learn deep patterns across all proteins, including where there is no experimental structure. There is evidence for local patterns in sequences that generalize beyond individual protein families, in the form of motifs that are local in the sequence (23) as well as motifs that are local in 3d space (24). However, the mapping between sequence and structure is not one-to-one (25), and designing sequences to reach a well-folded native state requires solving an exponentially large combinatorial problem to select a set of local sequence patterns which interact non-locally to specify a coherent structure (26). To design protein structure, the language model will have to develop an implicit understanding of how sequence determines structure, including local rules that link the design of structure with sequence, as well as global rules that determine whether a sequence is coherent and will fold into a native state.

### Generative protein design with language models

We evaluate language models generatively, focusing on generalization beyond natural proteins. The known protein sequences sampled by evolution represent only a small fraction of the vast number of possible proteins (Fig. 1A). To generalize outside the space of proteins that has been explored by evolution it will be necessary to access deep patterns of protein design that apply outside this space. We focus on two generative protein design tasks. The first is fixed backbone design where the objective is to generate a sequence that folds to the target structure. This task assesses the ability of the language model, which has been trained only on sequences, to design protein structures. The second task is free generation, where the structure is unconstrained and allowed to vary along with the sequence. This enables characterization of the full generative capability of the model across diverse sequences and structural patterns to understand the space of proteins accessible to the model.

**Figure 1.**
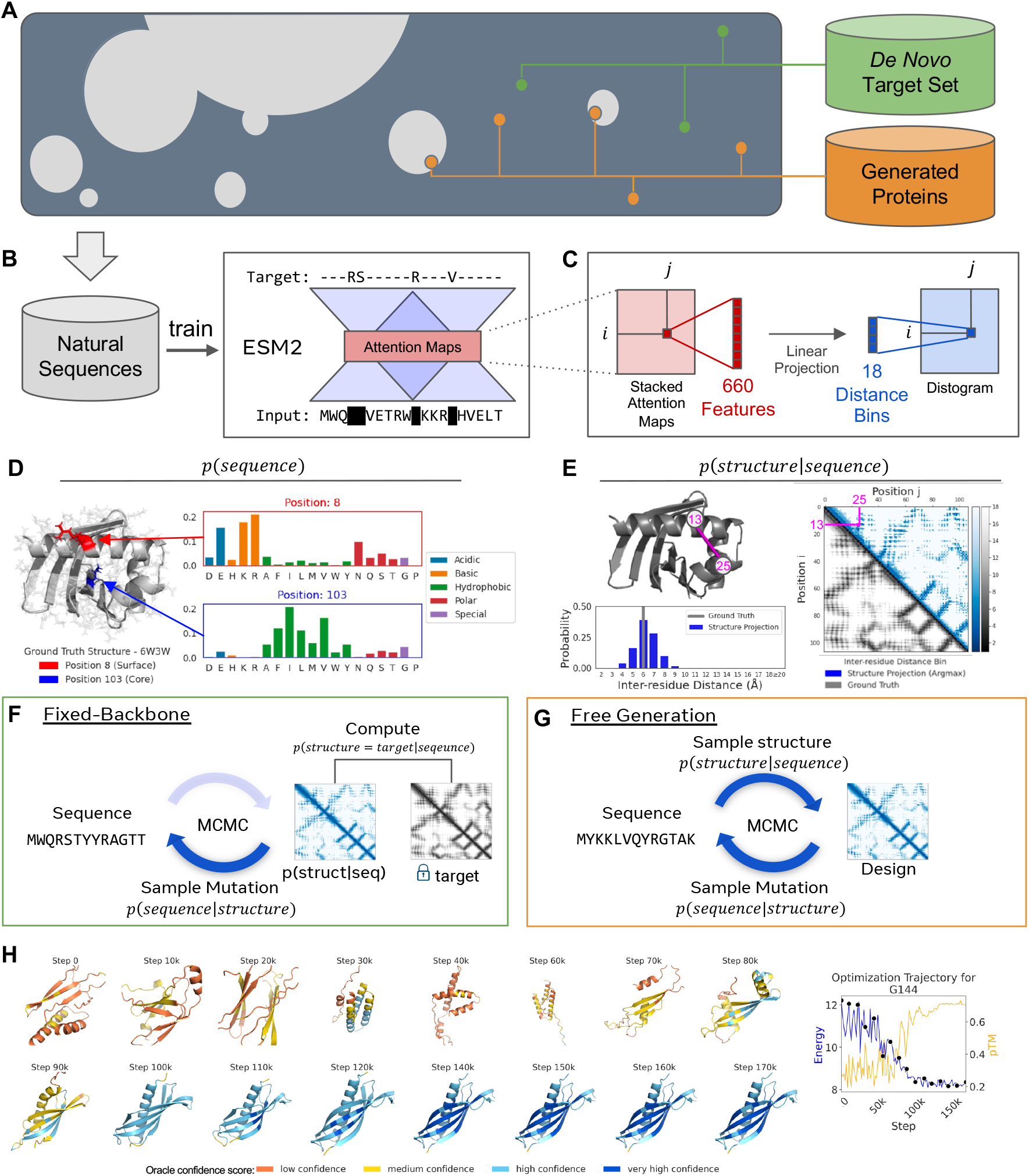
Overview. (**A**) Illustration of protein sequence space. Natural sequences (gray) cover a fraction of possible protein sequences. To generalize beyond natural sequences language models will need to access underlying design patterns. We evaluate language models on (i) a fixed backbone sequence design task with a set of *de novo* designed proteins (green), and (ii) via an unconstrained *de novo* protein generation task (orange). (**B**) The language model ESM2 is trained using masked language modeling over millions of diverse natural proteins across evolution. (**C**) After training, information about tertiary structure can be identified in the internal attention states of the model. A linear projection translates the attention at a pair of positions in the sequence to a distribution over inter-residue distances. (**D**) Probability of a sequence. The model outputs a probability for each amino acid at every position in the protein, here shown for the designed protein 6W3W. The model gives a higher probability to hydrophilic amino acids at a surface residue and hydrophobic ones at a residue in the core. (**E**) Probability of a structure given a sequence. For a given sequence the projection measures the compatibility of the internal representations of the language model with a structure. Tertiary structure is identified by probability mass on inter-residue distances less than 8Å. For 6W3W there is a good match between the projected structure (above diagonal) and ground truth structure (below diagonal). (**F**) The two terms giving the probability of sequences and structures are used to generate sequences. For fixed target design we use MCMC to generate sequences given a specified backbone structure, by sampling from the conditional distribution of sequences given a structure. (**G**) For unconstrained generation we allow both the sequence and structure to vary. (**H**) Predicted structures (using ÅlphaFold) are shown at even intervals across a single free generation trajectory. The model samples a range of possible topologies before narrowing to the refinement of one topology.

A test set of *de novo* designed artificial proteins is used to assess generalization beyond natural protein structures. The test set includes a diverse selection (N = 39) of structurally validated artificial protein structures from the Protein Data Bank (PDB) (27), which span a range of lengths (67 ≤ L ≤ 184), and topologies (Fig. S1 and Appendix A.1). Importantly, these *de novo* proteins have meaningful structural differences from proteins belonging to natural folds, including with respect to ideality, exact repetition, and symmetry of elements. Since the language model has not been trained on protein structures, generating designs for these backbones tests for the ability of the model to generalize to structures unlike those of the natural proteins whose sequences it has been trained on.

The language model, ESM2, is an evolutionary scale model of protein sequences that has been trained across the full extent of *natural* protein sequences (28). The training dataset excludes artificial sequences, as well as any sequences having similarity to the test set of *de novo* proteins used in the evaluations (Appendix A.1). ESM2 is trained with the masked language modeling objective (29) to recover the identity of amino acids from their context in the rest of the sequence (Fig. 1B). This training objective has been shown to materialize information about the folded structure of proteins in the internal representations of the model (7–9, 30). Since the training of the language model is only on sequences, information about structure that emerges must be the result of the unsupervised learning of patterns in sequences.

A linear projection from the attention maps of the language model identifies internal states that reflect protein structure. Previous work has shown that specific attention maps in transformer protein language models such as ESM2 encode the proximity of residue pairs in the structure (9, 30). We fit a linear projection that takes the attention between two positions in the protein sequence and outputs a distribution over pairwise distances (Fig. 1C). This maps an internal attention state of 660 dimensions into 18 inter-residue distance bins. Because of the limited number of parameters (660 per distance bin for a total of 11,898 including a bias for each distance bin), far too few to represent the immense complexity of possible protein structures, the output can be interpreted as a projection of the structure captured by the internal states of the model. The projection defines an energy landscape (a function of the representation states of the language model rather than a physical energy) that can be used to evaluate the compatibility of any given structure with the representation of the sequence produced by the language model. Application to the *de novo* target set shows understanding of existing *de novo* proteins (Table S1 and Figs. S2 and S3).

Together, the models of sequence, and structure given sequence, specify a generative model of proteins defined by the language model. The sequence model assigns a probability to any sequence, by giving a probability for each amino acid at every position in the protein (Fig. 1D). For natural proteins these probabilities reflect the functional effects of mutations, structural preferences of amino acids, and aspects of biochemical function (31). The projection of structure gives a compatibility between the language model’s representation of a sequence with a three dimensional structure (Fig. 1E). In this work, we consider these models to specify a generative model for protein design:

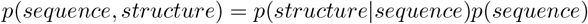

For fixed backbone design, protein sequences are generated by taking low temperature samples from the conditional distribution specified by the language model via Markov chain Monte Carlo (MCMC) with simulated annealing (Fig. 1F, Appendix A.3.1). Free generation removes the constraint on structure entirely and generates new proteins by sampling from the joint distribution of sequence and structure specified by the language model. A blocked Gibbs sampling approach is introduced which alternates between sampling a new structure conditioned on the current sequence, and sampling a new sequence conditioned on the current structure (Fig. 1G, Appendix A.3.3). An example free generation trajectory is shown in Fig. 1H. As the temperature is lowered, the trajectory proceeds from a phase where it samples a range of possible topologies before narrowing into a single topology that is refined into a confidently predicted structure in the final stage of optimization.

We perform extensive experimental testing of a total of 228 designs from the language model. Designs are considered a success that are well expressed, soluble, and pass a size exclusion chromatography (SEC) test for molecular (hydrodynamic) radius indicative of a properly-folded monomeric species (Appendix A.7). Experimental success of a significant fraction of the generated proteins, along with independent computational evaluation of the structures, demonstrates that language models are able to access a design space beyond that of natural proteins.

### Language models design sequences that fold to *de novo* structures

Fixed backbone design evaluates generation of sequences to realize a specified target structure. Use of *de novo* designed structures as targets requires the model to generalize beyond natural proteins, necessitating the use of more general patterns for the design of structure. Success at this task would indicate that the model has an understanding of the underlying design principles of protein structure generalizing to structures not encoded by natural sequences.

Across the test set of 39 artificially designed *de novo* protein structures, fixed backbone designs generated by the language model are predicted to closely match the target structures by the AlphaFold high-resolution structure prediction oracle. We generate 200 different designs for each of the *de novo* target structures (Appendix A.4). The generative model succeeds in producing low-RMSD designs for the vast majority of targets in the *de novo* test set (Fig. 2A). Subsetting to the best 10 of 200 designs by the language model’s optimization objective, median RMSD is < 2.5A for 84% (33/39) of targets and minimum RMSD is < 2A for 90% (35/39) of targets. Structures are also confidently predicted, with median pTM > 0.7 for 56% (22/39) and maximum pTM *>* 0.7 for 90% (35/39). Average sequence identity with the targets is low (22%), indicating that the language model is finding solutions to the design problem that differ from the original sequence.

**Figure 2.**
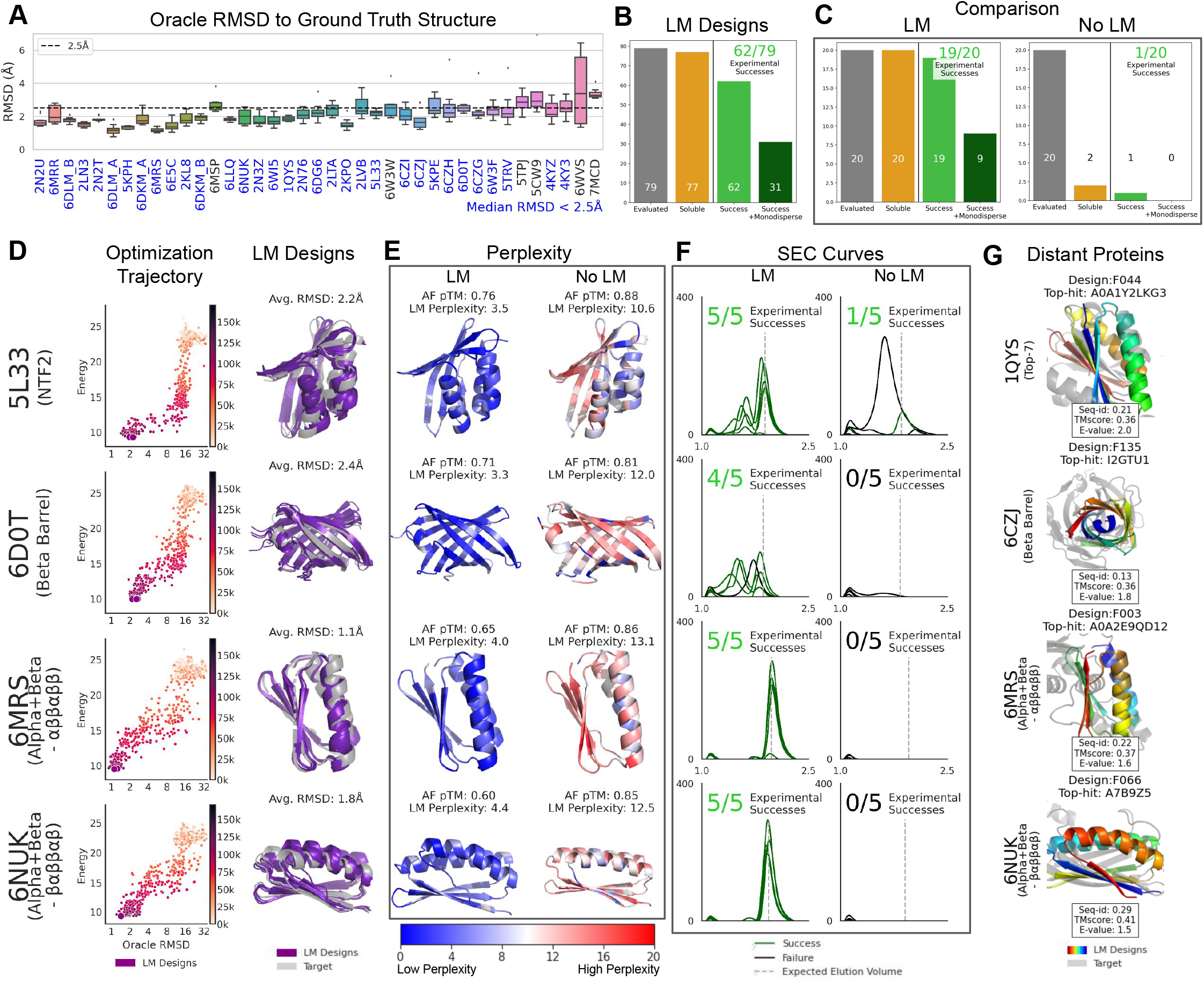
Design of sequences for de novo structures. (**A**) Overall evaluation of designs for the *de novo* target set using an *in silico* oracle. Rootmean-square deviation (RMSD) between C-alpha atoms designed structure (oracle prediction) and target structure is plotted for the top 10 designs by optimization objective for each target. Targets are ordered by increasing length. The language model generates sequences that are predicted to fold to the target structure for a large majority of *de novo* backbones in the test set. (33/39 achieve median RMSD < 2.5Å). (**B**) Experimental outcomes for ESM designs. A total of 79 designs across 6 *de novo* backbone targets were selected by a variety of criteria including sequence novelty and manual inspection for interesting motifs. Designs are considered a success if they are soluble and there is a peak at the expected elution volume by size-exclusion chromatography (SEC). Designs are categorized as monodisperse when the only peak is at the expected elution volume. Overall, 78% succeed, and 39% are monodisperse. (**C**) Experimental outcomes for comparison of designs with and without the language model. For each of the four targets, the top 5 out of 200 designs by optimization objective were selected for experimental evaluation. Overall 95% of designs with a language model succeed, while most designs without a language model fail due to insolubility. (**D**) (Left) Optimization trajectory showing energy specified by the language model vs RMSD to target, over the course of MCMC optimization. Energy decreases and funnels to low RMSD. (Right) Visualization of the top 5 designs selected by energy at the end of each trajectory. (**E**) Language modeling perplexity of designs. Language model designs are seen as probable by the language model, while high perplexity for the baseline designs indicates their sequences are seen as improbable. This coincides with experimental success. (**F**) Comparison of SEC traces between designs with and without a language model. The vast majority of language model designs are soluble and have a peak at the expected elution volume; in comparison few designs without a language model are soluble. (**G**) A subset of additional, successful language model designs are novel with respect to known natural proteins. Examples for four different backbones are shown with the design superimposed on the predicted structure of the top-significance hit from a sequence search against natural proteins. In each case the closest natural sequence has low sequence identity (<0.3) and predicted structure with different topology.

Generated proteins have high overall experimental success rates in the laboratory. We ran an additional set of fixed backbone design trajectories to explore the diversity of design motifs generated from the model. A total of 79 fixed backbone designs spanning 6 *de novo* targets were selected from a pool including the additional trajectories for evaluation by a variety of criteria including the presence of interesting structural motifs (Appendix A.6). Out of this set of experimentally tested proteins, 97% (77/79) were soluble, 78% (62/79) were successful, passing a SEC test for the presence of a peak at the expected elution volume indicating a folded monomeric species, and 39% (31/79) were monodisperse, exhibiting only a single SEC peak at the expected elution volume (Fig. 2B). Successes span a range of topologies, including a success for length 182 *de novo* TIM-barrel 6WVS which has a highly idealized symmetric structure (Fig. S4). Across the set of experimental successes, sequence identity to the original sequence of the target structure is low (mean = 24%), which suggests that the language model is exploring a new design space for the target structures.

We perform a controlled experiment to understand the role of the language model in experimental success of designs. For comparison we use AlphaFold as a model of the probability of structure given sequence. For a set of four fixed backbone *de novo* targets with distinct folds, we generate 200 designs using each method, with the top 5 by optimization objective for each method selected for experimental evaluation (Appendix A.3). Experimentally, 95% (19/20) of language model sequence designs and 5% (1/20) designs without a language model were successful (Fig. 2C). Augmenting AlphaFold with an n-gram prior, fails to rescue the designs (0% success rate, 0/20) (Tables S3 and S4).

Language model perplexity separates success from failure across both design methods. MCMC trajectories for the language model funnel to low RMSD with decreasing energy, with average RMSD values ranging from 1.1Å to 2.4Å (Fig. 2D). Notably, while ÅlphaFold confidently predicts the structures of language model designs, the language model does not assign high sequence likelihoods to ÅlphaFold designs. Language model perplexities of select ÅlphaFold-designed sequences range from 10.6 to 13.1 (Fig. 2E), significantly higher than the average *de novo* target sequence perplexity of 6.7. Other metrics have limited ability to identify experimental success (Fig. S5 and Table S4): the Rosetta all-atom energy function for modeling and design (32, 33) judges both sets to be good designs, packing metrics are similar but slightly favor the (unsuccessful) ÅlphaFold designs, while hydrophobicity and SÅP score favor the language model designs. Recently autoregressive inverse folding models directly conditioned on the target structure have demonstrated high experimental success rates in the laboratory (15). We generated sequences with ProteinMPNN and ESM-IF1 (14). Both models achieve high local confidence pLDDT (> 90 mean). Their ESM pseudo-perplexity is 5.76 and 5.79 respectively, higher than ESM designs and significantly lower than ÅlphaFold designs (Table S2), in line with high experimental success rates reported for those methods.

Experimental evaluation of both design sets (with and without the language model) indicates that 19/20 of language model designs are successful and 9/20 are monomeric (Fig. 2F). Target 6D0T has no monomeric designs from the language model, though the ground truth *de novo* sequence was also found to not be monomeric, when tested as a positive control (Åppendix Å.7). Designs without a language model largely fail due to insolubility.

Including the results of the controlled comparison, and the larger set of designs evaluated, the language model produced experimentally successful designs for all of a total of 8 *de novo* backbones. One possibility is that language model designs succeed because the model retrieves a protein similar to the target from its training set. To rule this out, we analyze the overall set of 81 experimental successes. Each design is searched against UniRef90 (which fully includes the sequences used to train the language model) to identify similar sequences (Åppendix Å.5). For 17 successful designs spanning 4 backbones, there are no significant (E-value < 1) sequence matches whatsoever in the training set. Four of these are shown in Fig. 2G. Of the remaining 64, sequence identity to the nearest sequence match is only 27% on average, and is < 30% for 41 of the 64, spanning each of the 8 tested backbones. This suggests that the model is not solving the design problem by retrieving similar sequences it has memorized.

To further understand whether the model is using homology at the threshold of detection by sequence similarity, we obtained AlphaFold predicted structures of hits, including those that do not reach the significance cutoff (Appendix A.5; Fig. S6). For 19/81 experimental successes top Jackhmmer hits are not structural matches to the design. For 19 designs spanning 4 backbones, the top-10 jackhmmer hits (including those that do not reach the significant threshold) all have TM-score < 0.6. For 8 of those designs spanning the same 4 backbones, top-10 hits are all likely to be a different fold (TMscore < 0.5). This suggests that while in some cases the model is able to use sequence homology at the threshold of detection, there are also cases where it appears to have generalized beyond that, further evidence that in many cases the language model is generating novel solutions to the design problem which differ both from the ground-truth sequence, and natural proteins.

### Language models materialize deep patterns of protein design

Generated proteins show evidence of using deep patterns of the design of protein structure. These patterns occur in the form of structural motifs used in the design of natural proteins applied in significantly differing sequence contexts, as well as the formation of motifs which cannot be found in related structures. Two well-studied ways that sequence determines structure are through amino acids that constrain backbone-geometry, and through the role of chemically diverse side chains in determining the intermolecular forces that stabilize a protein’s particular folded conformation. Two amino acids which influence backbone geometry are proline and glycine. These two amino acids add flexibility to and bend protein backbones, respectively. In three example designs, the language model places these residues to induce curvature in various secondary structure elements: a proline bends an alpha-helix, regular placement of glycines in beta-sheets promote the flexibility to form a beta-barrel, and all but one glycine are placed in loops in an NTF2 design (Fig. 3A). A side chain based motif present through fixed backbone designs is helix dipole capping, where side chains of amino acids at the ends of alpha-helices obscure otherwise exposed polar backbone atoms in the final alpha-helix turn (Fig. 3B). A second side chain based motif is hydrogen-bond networks in bulge-containing beta-turns, which are present in fixed backbone designs for beta-barrels, such as 6D0T and 6CZJ (Fig. 3C). This and to a larger extent the periodic glycines in beta-strands in Fig. 3A were identified as natural motifs that enabled successful *de novo* design of the target beta-barrel in (34).

**Figure 3.**
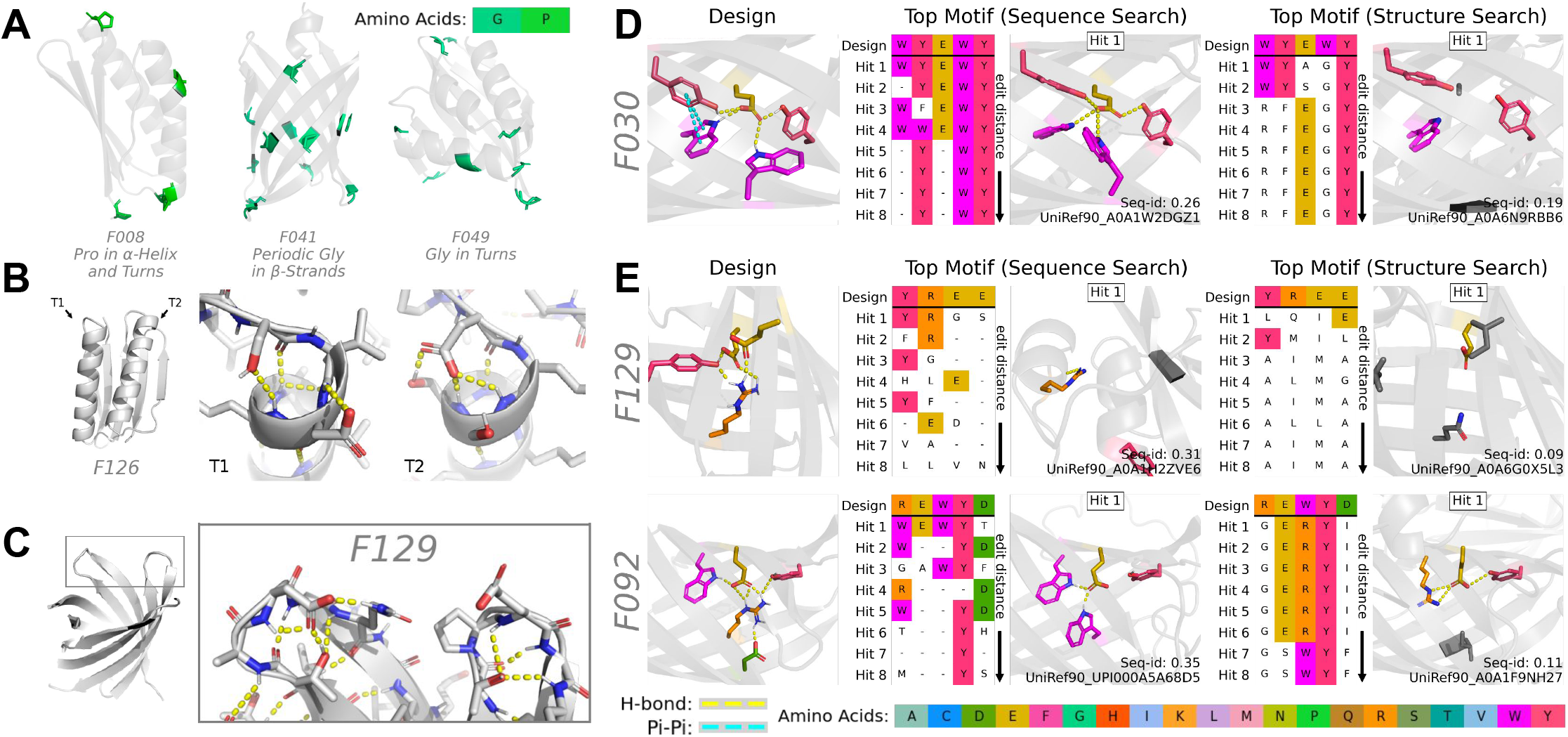
Language models materialize deep patterns of protein design, generating native-like and de novo motifs. (**A**) Placement of proline or glycine within three different designed proteins induce curvature on alpha-helices, beta-sheets, and turns. (**B,C**) Hydrogen bond networks in turns. (**B**) Helix dipole capping forms hydrogen bonds to obscure polar backbone atoms in the final helix turn. (**C**) Hydrogen bond networks formed in turns involving beta-sheets. (**D,E**) Comparison of motifs in designed and natural proteins. Designs (left) are compared against the nearest motif in natural proteins found by sequence search (center), and structure search (right). Hits are sorted by matching amino acids only at motif positions. (**D**) Example of a hydrogen bond motif used in one of the designs. Sequence matches are found that have the same motif in aligned positions. However the surrounding sequence context is significantly different, having 26% sequence identity. (**E**) Examples of possible *de novo* hydrogen-bond networks. Not only is the sequence context different, the motif itself is not present in the aligned positions of any matching natural sequences or structures.

Designs also exhibit complex hydrogen bonding networks. Some design successes include hydrogen bonding networks between four or more polar and even charged residues in the interior of structures. Design of buried polar and charged interactions is difficult due to the geometric constraints of energetically satisfying such interactions (35). Notably, the bond networks shown span a range of intermolecular force categories: among predicted structures, F129, a beta-barrel, contains a salt-bridge, F025 contains a pi-cation bond, and F030 contains a T-shaped pi-pi interaction (Fig. S7). The original designs for the examples shown have purely hydrophobic interiors. While these hydrogen bonding networks can only be fully confirmed by high-resolution structural studies, the biophysical properties observed (high yield of monodisperse protein with the expected retention volume) is consistent with their accuracy, since inaccurate placement of these residues is likely to lead to mis-folding and aggregation.

The hydrogen-bonding networks with polar residues are realized in new sequence contexts, indicating a strong form of generalization beyond the sequences used for training the model. We retrieve the most similar aligned sequences via Jackhmmer search of UniRef90 and similar, aligned structures via Foldseek (36) search of AlphaFold DB (37). Returned sequences are all sorted by minimum edit distance at aligned motif positions, and the closest matching motif is shown. (Appendix A.5.4). For generated protein F030 (Fig. 3D, Fig. S7), sequence search does recover a natural protein with this motif in aligned positions. However the surrounding sequence context in the design is dissimilar, having a full-sequence identity of 26%. For F129 and F092 (Fig. 3E, Fig. S7), not only does the surrounding sequence context have low sequence identity, the motif itself is not present in the aligned positions of any matching natural sequences or structures. Use of these motifs in fixed backbone designs is a remarkable form of generalization, since the model is applying them in new sequence contexts, and structures that are distinct from natural proteins.

### Language models generate novel structures and sequences

Language models generate new protein sequences that differ significantly from natural sequences. We sample a large set (N = 25,000) of proteins of fixed length (L = 100) without constraint on the structure. The blocked Gibbs sampling method which traverses the joint landscape of sequence and structure provides a more diverse set of proteins than previous unconstrained generation methods (Table S5).

Generations cover a variety of topologies with sequences overall dissimilar from natural proteins. Structures are predicted for all generated sequences using Alphafold, and generations are projected into two dimensions using t-SNE based on their pairwise structural distance measured by TM-score (Fig. 4A). In a hierarchical clustering of the structures, 7,663 distinct clusters were identified at a TM-score threshold of 0.75. The distribution of the generated secondary structures reveals a range of patterns with 52% of generations containing mostly alpha helices, 22% containing mostly beta sheets, and 28% a mix of alpha helices and beta sheets (Fig. 4B). A large fraction of the generations are well predicted by the oracle (median pLDDT = 84.49, 70% pLDDT > 70; Fig. 4C).

**Figure 4.**
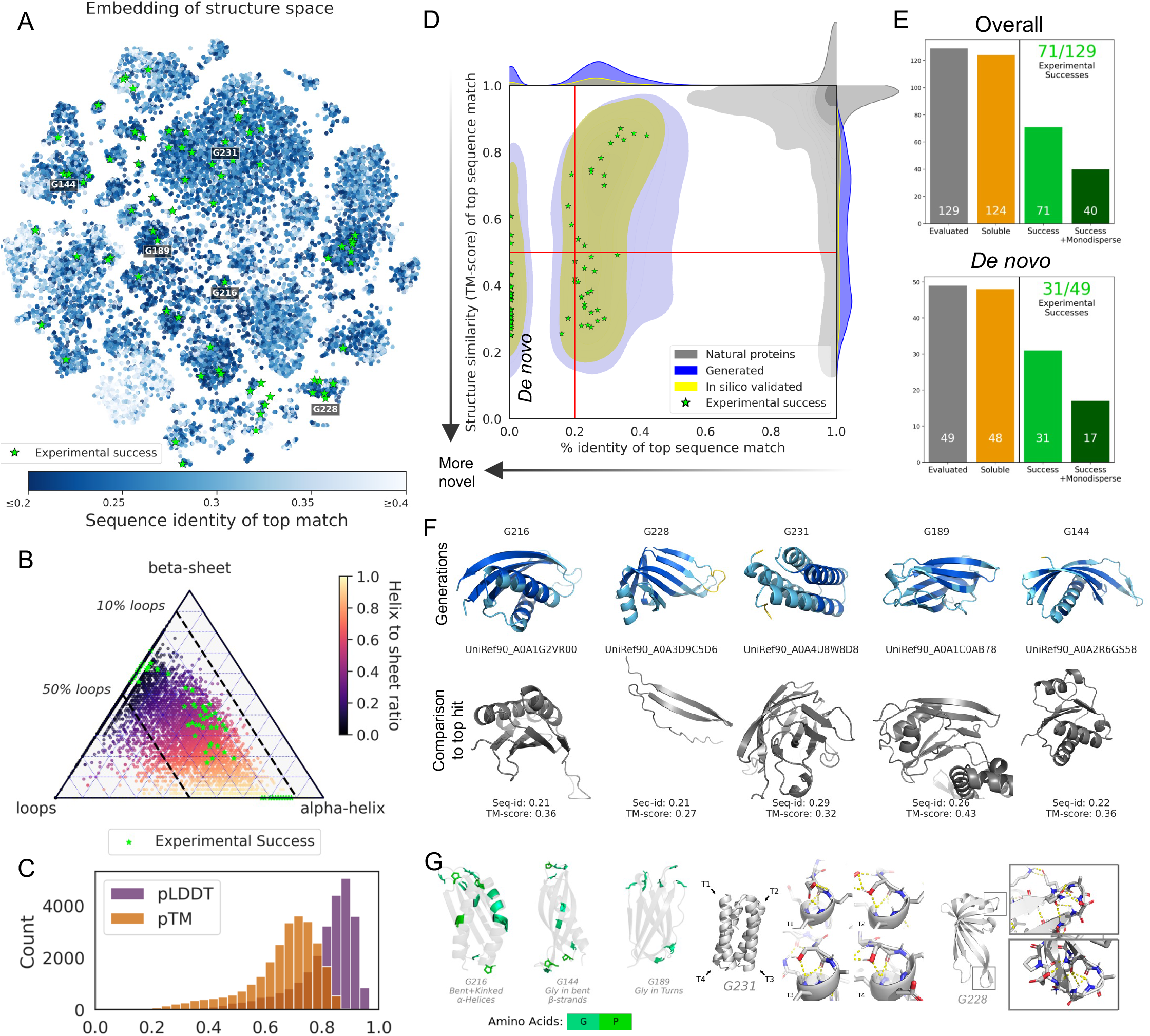
Language models generate novel structures and sequences. (**A**) Embedding of the structural space spanned by the generated proteins using t-SNE. Color indicates sequence identity to the best matching native sequence. A large fraction of the space has low sequence identity to natural proteins with 16% of generations having no significant sequence match to a natural protein. Designs that succeeded in experimental evaluation are indicated with green stars. (**B**) Distribution of secondary structure for generations. Experimental successes (green) are observed over differing compositions of secondary structure. (**C**) Distributions of pLDDT and pTM indicate designs are well predicted (median pLDDT of 84.5) by the *in silico* oracle. (**D**) Density plot of sequence and structural similarity to natural proteins. For each generated protein the best matching native sequence is retrieved from AlphaFoldDB. Each generated protein is plotted by its sequence similarity (x-axis) and structure similarity (y-axis) to the match, with hits that do not pass the significance threshold marked at zero on the x-axis. Generated proteins occupy a part of the space distinct from natural proteins, with a fraction having minimal sequence similarity to natural proteins (lower left quadrant). Designs passing *in silico* filters and experimental successes are coextensive with the overall distribution of generations. (**E**) Overall outcome of experimental evaluations. The majority of tested designs (55%) passed the solubility test and had an elution volume peak in the correct confidence interval (top). Additionally a high fraction (63%) of the evaluated proteins distant from natural sequences are successful (bottom). (**F**) Predicted structures of six experimental successes (top). Structures are aligned against the oracle predicted structure of their top significant hit from a sequence search of natural proteins (bottom); in all examples the predicted topology is different. (**G**) For generations in panel F, the same motifs as in Fig. 3A – 3C are shown: Proline and Glycine inducing curvature, helix capping, and hydrogen-bond networks in turns. Even on proteins with minimal similarity to naturals, the language model produces known motifs.

Many of the generations are distant in sequence from natural proteins. We measure the distance of generated sequences from natural proteins by searching each generation against the 200M natural sequences in AlphaFold DB (37). This also enables comparison of the structure of the nearest sequence match to that of the generated protein. Overall the language model generates proteins which show a clear separation from the distribution of natural proteins, including a fraction that are distant from known proteins. Fig. 4D shows the distribution of similarity to known proteins, where each generation is plotted according to its sequence (x-axis) and structural (y-axis) similarity to its top sequence hit, with insignificant (E-value > 1) hits placed at x=0 (16.6% of generations, in total). A large part of the distribution of generated proteins have structures different from those predicted for their nearest sequence match, further evidence that the model is not simply memorizing known proteins. A set of 15k natural proteins are also shown. Natural proteins cluster in the upper right corner, while generated proteins occupy a distinct part of the space. A significant fraction of the language model generated proteins (15.5%) have minimal similarity to natural proteins (lower left quadrant), with minimal sequence similarity (Seq-id < 0.2) of the nearest match, and a predicted structure likely to be a different fold (TM-score < 0.5).

A large fraction of the designs, including those that are distant from natural proteins, succeed experimentally. We selected a number of designs that passed our *in silico* quality filters for experimental evaluation. Out of the total set of generations, 20% (N = 5,198) passed the quality filters (Appendix A.4). A total of 129 of that set were expressed and evaluated, and 55% (71/129) were found to be experimentally successful. The 71 structures and their metrics are shown in Fig. S8, marked with a green star in Figs. 4A, 4B and 4D. Overall, 96% of the free generations that were evaluated were soluble, 55% had an elution volume peak in the correct confidence interval, and 30% were monodisperse (Fig. 4E top, Appendix A.7).

A high success rate is also observed for generations that are distant from natural proteins. For a set of 49 distant generations (Fig. 4D, bottom-left quadrant), 31 of 49 (63%) are successful in experimental evaluation. For these 31 experimental successes we perform a deeper analysis of similarity to natural proteins. We further search each against UniRef90 which provides comprehensive coverage of natural proteins and fully contains the language model’s training set. Out of 31 distant designs, 16 have no significant (E-value < 1) sequence matches whatsoever (Fig. S9). We obtain predicted structures for the top-10 sequence matches regardless of their significance. For 12 out of the 31 distant designs (5 of which are shown in Fig. 4F), none of the sequence matches are likely to have the same fold (TM-score < 0.5) (Fig. S9). Predicted structures are generally confident (78% of predictions with pLDDT > 70, average pLDDT = 81.24). Structural motifs observed in fixed backbone designs such as proline and glycine placement, helix capping, and hydrogen-bond networks, also appear within *de novo* generations (Fig. 4G). As a whole these results show that the language model generalizes outside the space of natural proteins to generate *de novo* proteins.

### Evolutionary scale language models

Transformer protein language models were introduced by (7), which found evidence for the emergence of information about function and tertiary structure from the unsupervised training. Concurrent work at a smaller scale examined LSTM-based models (38–40). Large scale protein language models with billions of parameters have now been open sourced (8, 41–43). Generative use of language models has recently been explored by *in silico* studies (44, 45), and experimentally with confirmation of function for new sequences generated for existing protein families (22). To the best of our knowledge, experimentally validated work (20, 22, 46) with sequence based models has not crossed the threshold of < 30% identity to natural proteins.

## Conclusions

The classical picture of sequence space as being constituted by independent local evolutionary landscapes around each protein family would suggest that language models will be limited to a memorization of the space of natural proteins. Consistent with this, the information about structure that emerges in language models of proteins has been shown to depend on the evolutionary information available to the model during training, which would appear to be unencouraging for the potential to use language models generatively beyond natural proteins. Here we have presented evidence counter to this: language models generalize beyond natural protein families to generate proteins in a sequence space distant from natural proteins. Our results are the first time purely sequence based approaches have been shown to generalize beyond natural proteins, and are promising for sequence based generative artificial intelligence for *de novo* protein design, where we have demonstrated that there exists a space of *de novo* proteins, distant from those in nature, that are designable by generative language models.

This generalization points to a deeper structure underlying natural sequences, and to the existence of a deep grammar that is learnable by a language model. Our results suggest that the vast extent of protein sequences created through evolution contains an image of biological structure and function that reveals design patterns that apply across proteins, that can be learned and recombined by a fully sequence based model. The generalization beyond natural proteins does not necessarily indicate that language models are learning a physical energy. Language models may still be learning patterns, rather than the physical energy, but speculatively, in the limit of infinite sequence data, these patterns might approximate the physical energy. At a minimum the language model must have developed an understanding of the global coherence of a protein connecting the sequence and folded structure.

The existence of a deep grammar across proteins would explain the two observations which prima facie seem to contradict each other: that the understanding of natural proteins depends on evolutionary support in the training data, and also that the language models generalize outside of known natural protein families. If there is a power law distribution of learnable patterns, then it is expected that many protein structures will be designable with the common patterns that have the most support in the training data. At the same time, the frequency that patterns are observed in the training data will correspond with the learnability of the patterns. It will take greater amounts of training data, and model capacity, to learn rare patterns. This is consistent with the observation of both generalization to a new design space (that is accessible via the patterns that have been learned), and dependence on support in training data (the proteins composed of rare patterns are harder to learn). If scaling laws continue to hold for protein language models we can expect their generative ability will continue to improve. As models and data scale, the existence of a learnable underlying grammar would predict that the rare patterns will be learned, expanding both the predictive ability of the model, and the design space that is accessible.

## Acknowledgements

We would like to thank Halil Åkin, Salvatore Candido, Brian Hie, Ådam Lerer, Zeming Lin, Wenting Lu, Roshan Rao, Yaniv Shmueli, Nikita Smetanin, and Zhongkai Zhu for technical help, feedback, and discussions that helped shape this project. We thank Christoffer Norn and Ånastassia Vorobieva for insights into *de novo* protein structures, and Ivan Anishchenko for sharing the *de novo* protein target set. We thank Gabriel Rocklin and Kotaro Tsuboyama for stability experiments when the project was nascent. We thank Laurens van der Maaten, Ammar Rizvi, Jon Shepard, and Joe Spisak for program support.

This work was supported with funds provided by the Audacious Project at the Institute for Protein Design (D.B.), an EMBO longterm fellowship (ALTF 139-2018, to B.I.M.W.), the Open Philanthropy Project Improving Protein Design Fund (J.D., and D.B.), a Human Frontier Science Program Cross Disciplinary Fellowship (LT000395/2020-C, to L.F.M.), an EMBO Non-Stipendiary Fellowship (ALTF 1047-2019, to L.F.M.), a gift from Meta (D.B.), and the Howard Hughes Medical Institute (D.B.). For this project, S.O. is supported by NIH Grant DP5OD026389 and NSF Grant MCB2032259.

## A. Methods

### A.1. Data

#### A. 1.1. *De Novo* Target Set

A held-out set of *de novo* proteins is used for the task of design with a fixed target backbone. A diverse set (*N* = 39) of *de novo* structures from the Protein Data Bank (27) was selected, which span a range of lengths (67 ≤ *L* ≤ 184), folds (e.g. alpha-bundle, beta-barrel, NTF2, Rossman) and *de novo* design methods (26, 34,47–56). See Fig. S1 for a visual display of all x-ray crystal structures comprising the *de novo* target set. These proteins were designed by humans, rather than by natural evolutionary processes. Importantly, these *de novo* proteins have meaningful structural differences from proteins belonging to natural folds. For example, NTF2 targets have unnatural binding pockets (55), beta-barrels are narrower and have short beta-turns (34), and some designs were entirely new folds at the time of their creation (47). Although these proteins are by definition distinct in both sequence and structure from natural proteins, each protein in the target set is queried against UniRef100 (28), which subsumes the training set of ESM2, and all sequences returned as matches by Jackhmmer search are excluded from the language model’s training, see next section.

The Protein Data Bank Identification Codes (PDB IDs) of the *de novo* target set are:

1QYS,2KL8,2KPO,2LN3,2LTA,2LVB,2N2T,2N2U,2N3Z 2N76,4KY3,4KYZ,5CW9,5KPE,5KPH,5L33,5TPJ,5TRV 6CZG,6CZH,6CZI,6CZJ,6D0T,6DG6,6DKM_A,6DKM_B, 6DLM_A,6DLM_B,6E5C,6LLQ,6MRR,6MRS,6MSP,6NUK, 6W3F,6W3W,6WI5,6WVS,7MCD

#### A. 1.2. Sequence dataset used to train esm2

The language model used throughout this work is ESM2_650M (8). Therefore, all pretraining settings described in that work apply for the language model used here.

To test whether the language model’s understanding of proteins generalizes from natural to *de novo* space, it is critical that the model did not see *de novo* proteins at train time. To this end, we first remove all sequences from ESM2’s train set labeled as “artificial construct” on the UniProt (57) website, when 2021_04 was the most recent release (1,027 total proteins). To guard against mislabeled proteins, and to further remove sequences in the train set which may bear similarity to the target set, we additionally perform Jackhmmer (58) searches of each *de novo* sequence against UniRef100 `2021_04` with flags `—num-iter` 1 {`max`, and remove all hits returned by the tool from ESM2’s training set (58,462 proteins).

#### A. 1.3. Structure projection dataset

The structure projection network was trained on a nonredundant dataset from PDB consisting of 15,051 proteins (structure release dates prior to 1 May 2018) used in Yang, et. al. (59).

#### A. 1.4. Heldout SET OF NATURAL PROTEINS

A small (*N* = 214) set of natural proteins with structures in the PDB was selected to serve as a baseline comparison when evaluating language model *de novo* protein understanding in Figs. S1 and S2 and Table S1. The set is composed of PDBs available on July 2020 that have sequence identity < 0.3 to the dataset used to train the structure projection, according to mmseqs2 (60). A length filter of 50 ≤ L < 250 was applied to roughly match the length distribution of the *de novo* target set (67 ≤ L ≤ 184).

### A.2. Models

#### A.2.1. ESM2

We use `ESM2_650M` (8) as our choice of large-scale protein language model throughout this work. ESM2 is a Transformer model trained via masked language modeling over the universe of known, natural protein sequences. At training time, protein sequences are shown to the model with a fraction of their residues masked, randomly permuted to a different amino acid, or left unmodified, according to standard BERT noise probabilities (7). The model’s task is to predict those masked residues given bi-directional context of all unmasked residues in the input. ESM2 is trained only on natural protein sequences; sequences annotated as artificially constructed and sequences matched by sequence search with *de novo* target set queries were removed from the language model’s training set Appendix A.1.2.

The language model is used to approximate *p*(*sequence*) via the pseudo-likelihood (61). Let us first define the probability *pθ* (*x_i_*|*x_-i_*) over the possible amino acids at position *i* in sequence *x*, conditioned on the remainder of sequence. This conditional probability is obtained by constructing *x*_-i_ where amino acid *i* is replaced with <`mask`>, and computing the language model probabilities at position *i*. The pseudo-likelihood is then defined as Π_*i*_ *p*(*x_i_*|*x_-i_*).

#### A.2.2. Structure Projection

The structure projection is a single learned affine projection (linear projection with bias term) from ESM2 internal representations to inter-residue distance, applied identically to each position-pair [*i*, *j*] of the protein.

In its implementation, the (*N* = 660) attention maps computed during ESM2 inference for a given sequence are used as input to a linear projection. At position [*i*, *j*] we compute *z_ij_* with (*N* = 18) dimensions: *z_ij_* = *W_projection_*attention_maps_*ij*_ + *b_projection_*. The vectors *z_ij_* are the softmax logits which define the categorical distribution *p*(*d_ij_*|*sequence*) over binned inter-residue distance between the carbon-beta atoms, known as distogram (62). Under a conditional pairwise independence assumption we use Π_*i,j,j ╪*_ *P*(*d_ij_*|*sequence*) to approximate*p*(*structure|sequence*). There are 660 * 18 + 18 = 11, 898 total learned parameters in the structure projection. The binning resolution of the model is ≈1A, with 16 bins spanning the range [2.5A, 20A). The very first bin represents <2.5A, and the very last bin represents >20A. Symmetry was applied to prediction logits, since distograms are by definition symmetric (*d_ij_* = *d_ij_*). Weights of the ESM2 model were frozen during training of the structure projection.

The structure projection was trained on a random subset of 80% of the sequence and structure pairs published in Yang, et. al (59) Appendix A.1.3. As in that work, distograms are constructed from inferred Carbon-beta coordinates of protein backbones. We trained the model for 10 epochs with a batch size of 4 and learning rate of 1e-2 using categorical cross-entropy loss between all [*i, j*] pairs in the predicted distogram and ground truth distogram. There are no common structures between the dataset used to learn the structure projection and the *de novo* target set.

#### A.2.3. N-gram Prior

Background distributions of uni-, bi-, and tri-gram (n-gram) amino acid frequencies were determined via the amino acid frequencies in UniRef50, release `2018_03`. During design, the Kullback-Leibler divergence (*D_KL_*) is calculated between the n-gram frequencies of the background distribution and of the designed sequence. *D_KL_* terms are added with equal weight to produce a single n-gram energy term. Conceptually this can also be seen as using the n-grams as a language model *p*(*sequence*) (63) which can be combined with the ESM transformer language model. Concretely, the energy function is defined between n-gram frequencies of our design sequence and background:

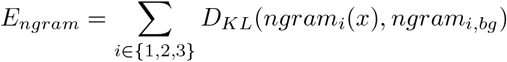

### A.3. Tasks

#### A.3.1. Fixed backbone design

The goal of fixed backbone design is to generate a protein sequence x for a target backbone y. As in (59), the backbone is derived from the set of 3D coordinates of the protein’s Carbon-beta atoms (inferred for glycines), with length equal to the number of residues in the protein. These 3D coordinates are converted to a distogram of binned pairwise distances Appendix A.2.2.

We’d like to sample sequences x with high likelihood, conditioned on the target backbone, Y:

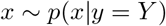

To sample from this distribution, we first note from Bayes rule that this is equivalent to sampling from the unnormalized product of unconditional sequence prior *p*(*x*) and a conditional structure distribution *p*(*y*|*x*):

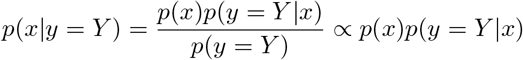

- *p*(*x*): approximated by the language model’s pseudolikelihood computed by multiplying marginal likelihoods when masking out each individual token, and the n-gram prior.
- *p*(*y* = *Y*|*x*): approximated by the distogram distribution from the language model with structure projection head, evaluated for the target *Y*.
- *p*(*y* = *Y*): constant, in the case of a fixed target.

To sample *p*(*x*|*y* = *Y*), we utilize an energy-based sampling procedure, via Markov-Chain Monte-Carlo (MCMC). Formally, our full energy function for sampling from *p*(*x*)*p*(*y* = *Y*|*x*) is the following expression for a randomly selected sequence index *i*:

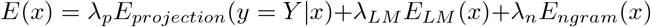

where:

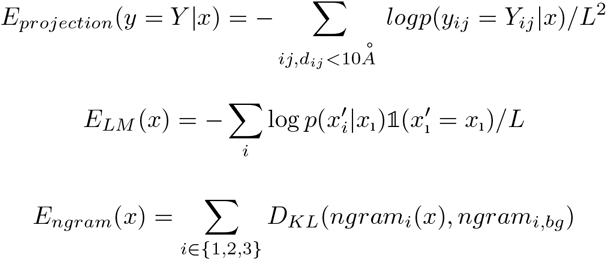

In the above energy function *E*(*x*), the first term *E_projection_* specifies sequence-structure consistency, taking only pair-positions into account which are in contact in the target, i.e. have inter-residue distance *d_ij_* < 10Å. The term *E_LM_* specifies sequence negative log likelihood, and the term *E_ngram_* is based on the n-gram model of sequence likelihood. The terms are composed together with separate weights *λ_p_* = 3, *λ_LM_* = 2, *λ_n_* = 1 enforcing different prioritizations of each objective, which were determined by hyperparameter sweeps. The overall energy function *E*(*x*) defines a Boltzmann distribution:

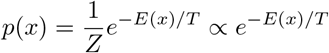

A uniform random amino acid mutation 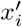 at a randomly selected sequence index *i* is proposed at each step with Metropolis acceptance rate *α*:

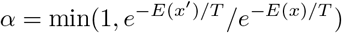

Mutations to cysteine were disallowed, as their presence would interfere with experimental evaluation. Note that by defining acceptance as the ratio of *E*(*x’*) and *E*(*x*), the relative ratio between *E_LM_* (*x’*) and *E_LM_* (*x*) can be efficiently approximated in 1 forward pass through the language model (by computing marginal substitution likelihoods at the substituted position), as opposed to the L forward passes required to explicitly compute *E_LM_*(*x*).

We perform 170,000 steps of MCMC sampling. We use a geometrically decaying temperature schedule for simulated annealing. Every 10,000 steps, we decay the temperature *T* by 2, from an initial value of 8 to a terminal value of approximately 6e-5. Full design trajectories take ≈ 10 hours for a fixed backbone design with sequence length ≈ 100 on a single 32GB Volta gpu. We could achieve successful designs on most targets (low target RMSD according to the oracle) with fewer steps, but the step count was increased to achieve best performance, especially for longer fixed backbone designs, e.g. 6WVS (*L* = 182).

#### A.3.2. Fixed BACKBONE DESIGN WITHOUT A LANGUAGE MODEL

Designs from the language model (“LM”) were compared against designs from a baseline with a powerful structure predictor, but no language model (“no-LM”). For this baseline, AlphaFold was used as the structure model.

To keep the comparison with LM designs matched, AlphaFold’s pairwise distance (distogram) output was used as *p*(*structure* = *Target|sequence*). Since no transformer language model is used, there is no *p*(*sequence*) term and fixed backbone design without a language model optimizes for sequences that have high likelihood *p*(*structure|sequence*). Additionally, to ensure a fully matched comparison against the LM designs, a second set of no-LM designs were generated, which include the same *E_ngram_* term used for LM designs Appendix A.2.3. The additional n-gram term can be interpreted as adding a weak n-gram language model. The coefficient of this n-gram term was selected via a line sweep Table S3. In the main comparison, we only feature results from the set without the n-gram term, since that set was more successful experimentally (1/20 successes vs. 0/20 successes with n-gram term).

A gradient-based public algorithm for producing AlphaFold-based designs was used. Baseline designs were produced by ColabDesign (12, 16) (commit hash e7bb3def), using the `design_3stage()` AfDesign recipe, which alternatingly and then simultaneously optimizes across all 5 AlphaFold pTM model replicas. It was found that more steps improved the convergence to low target RMSD over the course of optimization, so the default number of steps used was scaled up by a factor of 5, for a total of 1500 sofL·itcrs, 500 temp_iters, and 50 hard_iters. This design protocol requires less steps of optimization due to employing gradient-based optimization; the algorithm can update each position in the sequence at each step, whereas the MCMC protocol we employ only makes a single mutation at each step. Although AlphaFold’s distogram output was optimized rather than its atomic structure prediction output, all designs were verified to have < 1*Ä* RMSD to target and > 0.8 pTM according to the AlphaFold Oracle Appendix A.4.1.

LM and no-LM protocols were used to produce 200 designs per target each. Simple selection of the top 5/200 seeds (per target) according to each protocol’s optimization objective was used to select designs for experimental evaluation Appendix A.6.2.

#### A.3.3. Free GENERATION

The goal of free (unconstrained) generation is to design a new protein sequence x which is sampled from the universe of possible sequences x and their associated backbones y. As in the previous section, backbones of designs are represented by the distogram distribution over pairwise distances. In particular, we wish to sample sequences x and associated structures y with high joint probability:

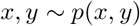

We utilize an energy-based sampling procedure to sample both sequence x and structure backbone y from this joint distribution. In particular, we utilize a blocked Gibbs MCMC sampling procedure where, starting from an initially random sequence x, we sample a definite backbone y for the current sequence

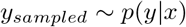

and then sample an updated sequence x’ given the current backbone y.

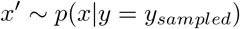

During the *p*(*y*|*x*) sampling phase, inter-residue distances are sampled independently at all pair-positions in the distogram. During the *p*(*x*|*y*) sampling phase, 3 steps of the MCMC protocol for fixed backbone design are performed (see prior section), where the sampled backbone *y_sampled_* is used as a target.

In total, 170,000 steps of MCMC are performed, where a step is comprised of a *p*(*y*|*x*) sampling phase and a *p*(*x*|*y*) sampling phase. For *p*(*x*|*y*) sampling, the same temperature schedule is used as in fixed backbone design: temperature is decayed by a factor of 2 every 10,000 steps, from 8 to ≈ 6e - 5. For the structure sampling step, a fixed temperature of 1 is used. Annealing both temperatures led to low diversity (alpha-bundle) solutions which indeed have very high *p*(*y*|*x*) and *p*(*x*|*y*), which did not happen with fixed *p*(*y*|*x*) temperature. With this protocol to sample from the joint distribution, a diverse set of topologies was generated with varied secondary structure content (Fig. 4A; Fig. 4B), respectively. Finally, as in fixed backbone design, mutations to cysteine were disallowed, as their presence would interfere with experimental testing.

### A.4. *In silico* Quality Metrics

#### A.4.1. Structure Oracle

Designed sequences were given as input to AlphaFold for an *in silico* assessment of their structure. AlphaFold serves as a powerful orthogonal predictor of protein structure, as the AlphaFold model differs from ESM2 in its architecture, objective, and training data. When evaluation designs, sequences are input without generating a multiple sequence alignment (MSA) nor using any templates. We follow the standard protocol of predicting structure and confidence scores (pTM captures global confidence, pLDDT captures local confidence) with all 5 publicly released models, then select the most confident output by pLDDT. Amber relaxation was performed on the selected, predicted structure. All predicted structures of designs in this study come from this pipeline, and structural metrics described in the following sections are calculated using these predictions. Confidence metrics pTM and pLDDT, as well as RMSD to target structure where available, can be used for selecting designs as well Appendix A.6.

#### A.4.2. Solubility AND AGGREGATION METRICS

Three *in silico* metrics are used for the purpose of filtering out candidates with strong evidence that they will not be soluble or monomeric:

1. Hydrophobic Solvent Accessible Surface Area (SASA) computes the SASA for each hydrophobic residue. It quantifies how much of the protein’s surface, which is accessible to the solvent, is hydrophobic. High hydrophobic SASA is problematic as for monomeric proteins we expect the SASA to be mostly polar in order to stay in solution rather than aggregating via exposed hydrophobic surface area.
2. Net Charge: a simplified sequence-based net charge by counting positively and negatively charged amino acids in the sequence, to try to avoid proteins with zero net charge, as this may lead to aggregation in polar solvents.
3. The spatial aggregation propensity (SAP) is a metric introduced to quantify the aggregation propensity, i.e. whether the protein will aggregate into non-functional and typically insoluble assemblies (64). SAP score identifies surface exposed patches of hydrophobic residues and computes a weighted sum of atoms with positive hydrophobicity score. The SAP metric is useful to study and filter both fixed backbone designs and free generations, as in both cases we aim to design monomeric proteins. This requires the monomer to be soluble and have mostly hydrophilic amino acids on the surface, corresponding to low SAP score (65). This metric is correlated but can be complementary to the hydrophobic SASA. For comparison across sequence lengths, we use average SAP score, i.e. averaged over residues.

For fixed backbone designs tested experimentally, we use SAP score during selection except for those in the “LM vs. no-LM” comparison where no additional in-silico metrics are used for filtering (Methods; Selection of designs for experimental evaluation). When filtering free generations, we use all three hydrophobicity metrics with relatively loose thresholds, and combine them with logical “and”, i.e. the candidate has to pass all filters. Firstly, hydrophobic SASA < 1.7 times the “ideal surface” computed using the ideal sphere for the same length protein. Secondly, we require sequence-based net charge ≥ 2 or ≤ −2. Finally, we filter for averaged SAP scores ≤ 0.4, and relax this threshold to 0.5 when the predicted structure contains at least 25% beta strands.

#### A.4.3. Packing Metrics

Two metrics are used to filter candidates which are likely not well-packed:

1. Protein Packing is quantified with the Rosetta PackStat filter, and is an approximate implementation of RosettaHoles (66). This is a stochastic algorithm, so it is averaged across 100 repeats. It returns a score between 0 and 1, where 1 means perfect packing. We keep free generation candidates only if packing > 0.55.
2. Shape Complementarity of secondary structure elements in the structure (67) is implemented in the Rosetta SSShapeComplementarity filter with loops=“true” helices=“true”. This metric aims to quantify whether the surface normals from different interacting secondary structures are well-aligned, indicating that secondary structure elements fit well together. We keep free generation candidates only if shape complementarity > 0. 6.

The Packing and Shape Complementary metrics were computed twice: once on the structure from the AlphaFold pipeline after Amber relaxation, and once after an additional step of Rosetta minimization with the `beta_nov16` (32). Logical “or” between structure filters is used: if either of the structures passes the filter, the filter is satisfied.

#### A.4.4. Globularity METRICS

A final set of metrics are used to screen out proteins which are not globular and have oblong shapes such as extended helix bundles. We follow Dill et al. (68) and define the idealized radius of a protein based on its number of residues as 2.24 * (numresidues ** 0.392) (68), and its corresponding ideal surface area based on this radius. Using these as reference values, we define relative SASA and relative radius of gyration. The following metrics and thresholds are used:

1. The Radius of Gyration is the root mean square distance from the center of mass (not taking residue weights into account). We keep candidates if the relative radius of gyration is < 1.5.
2. Total solvent accessible surface area (SASA) computed by Rosetta TotalSasa. We keep candidates if the relative SASA is < 3.
3. Contact Order was computed but we did not filter on this metric. The sample of free generations spanned a range of Contact Order values.

### A.5. Comparison to natural proteins

#### A.5.1. Settings USED FOR SEQUENCE SEARCH

Designed sequences are tested for their distance from natural protein sequences via querying them against large-scale sequence databases. We emphasize that comparisons described throughout this section (except in the case of motifs) are made for hits returned by sequence search; comparison of predicted structure for designs to known structure databases always returns hits likely to possess a similar fold (Fig. S10).

For sequence search, we use jackhmmer 3.3.2, a sequence search tool from the HMMER suite (58). Two jackhmmer settings were modified from their defaults, based on failure modes observed during during our analysis, which queries distant /de novo sequences against large-scale (> 100M sequences) search databases:

1. **One Iteration.** Jackhmmer was run with only 1 iteration, instead of multiple (the default). This change was made because it was observed that additional iterations resulted in a growing amount of returned spurious hits, when given distant sequences as input, such as de novo ground truth sequences. Specifically, for query sequences with few natural sequence homologs, false positives increasingly dominated the query profile used on subsequent jackhmmer search iterations.
2. **Sorting by E-value of the best-scoring domain.** Throughout this paper, Jackhmmer results are always sorted according to *best-domain* (rather than full-sequence) E-value. It was found that ranking hits by full-sequence E-value frequently led to more spurious top hits. Specifically, designs—which comprise a single domain in fixed backbone design, and have a single-domain-like globular structure in free generation—tended to match long, repetitive hit sequences containing a repeated domain. In these cases, multiple weak per-domain matches resulted in a high cumulative full-sequence significance, even though the design had no strong match to any single domain in the hit sequence. This is a known potential failure mode stated in the user’s guide (69). Though significance is determined by E-value for the best domain match, when top jackhmmer hits are subsequently analyzed (e.g. for calculating sequence identity and predicting structure) the *full* hit sequence is used.

Overall, Jackhmmer was run with non-default settings (-n 1 -seed 0).

Designs are compared against their sequence hits on three major axes: **E-value**, **Sequence Identity**, and **TM-score**:

1. **E-value.** Jackhmmer returns an E-value for each hit, which quantifies the significance of each hit’s sequence match to the query. Specifically, E-value is the number of false positives that are expected to score as or more strongly than the given hit due to random chance. Hits with a (best-domain) E-value < 1 are considered significant. At this significance level, we expect one hit on average to be falsely considered significant, when querying each design against some large sequence database.
2. **Sequence Identity.** Sequence identity of the design to each of its hits was calculated via local alignment with Biotite’s (70) `biotite.sequence.align.align_optimal()` given the BLOSUM62 substitution matrix applied to the full sequence of the design and the full sequence of the hit. Sequence identity was calculated as the number of matching characters in the two aligned sequences divided by the full length of the original query sequence (rather than just the length of the aligned region).
3. **TM-score.** Designs were also compared to some fraction of their top hits via TM-score of their predicted structures from the TM-align tool (71). Predicted structures of designs are obtained using the (AlphaFold, single-sequence) structure oracle (Appendix A.4.1). Predicted structures for (the full sequences of) top jackhmmer hits are obtained from AlphaFold DB, or the structure oracle given an MSA (instead of a single sequence) as input.

#### A.5.2. Comparison OF ALL FREE GENERATIONS TO NATURAL PROTEINS IN ALPHA Fold DB

In the case of Fig. 4D, each of the 25k free generations and the ≈15k natural proteins from (59) was queried against the sequences in AlphaFold DB (37), which comprise UniProt `2021_04` (57). Because all sequences in this database have a structure predicted by AlphaFold, searching against this database enables comparison of predicted structure at scale. We compare designs to only their single most significant (by best-domain E-value) hit, on the bases of sequence-identity and TM-score of predicted structures, fetched from the url: https://alphafold.ebi.ac.uk/files/AF-<UniProtID>-F1-model_v3.pdb. The bottom-left quadrant of Fig. 4D, where sequence-identity < 0.2 and TM-score of predicted structure < 0.5 was used to define a set of 49 distant free generations, of which 31 (67%) succeed experimentally. Generations that have no significant (best-domain E-value < 1) hits are displayed at 0 sequence identity in that plot, to distinguish them from generations possessing significant hits, visually. Results from this comparison are used in Fig. 4D, the definition of “49 distant free generations” in the Introduction, and the analysis of free generations.

#### A.5.3. Comparison OF EXPERIMENTALLY EVALUATED DESIGNS TO NATURAL PROTEINS IN UniRef90

In all other cases, when we compare experimentally evaluated designs to known natural proteins, we query against UniRef90 2021_04 (28), which fully contains the set of sequences seen by the language model during training.

Unlike in the comparison of designs natural proteins AlphaFold DB, where we consider top-hit statistics, only, we perform a more comprehensive analysis:

1. **E-value**. Same as the comparison to AlphaFold DB. (Best-domain) E-value of the top hit = *minimum* over *all* hits due to sorting.
2. **Sequence Identity.** Is calculated as a *maximum* over sequence identities for *all* significant (best domain E-value < 1) hits.
3. **TM-score.** Is calculated as a maximum over the top-10 sequence hits. Predicted structures are acquired https://alphafold.ebi.ac.uk/files/AF-<UniProtID>-F1-model_v3.pdb where possible. However, a fraction of UniRef90 proteins (≈ *20*%) are not present in AlphaFold DB. For these proteins, predicted structures were obtained by folding their (full) sequences via the structure oracle, given an MSA produced by jackhmmer on UniRef90 (same settings as in Jumper et al. (72) as input rather than just the single hit sequence. Of the 228*10=2280 total jackhmmer hits considered throughout this paper, 4 (1 significant) had their TM-scores omitted from analysis, due to not being in AlphaFold DB and failing during oracle structure prediction because of GPU memory limitations (all are length *> 1000*). These errors do not affect the sequence statistics of jackhmmer hits (sequence identities, E-values).

UniRef90 is not the exact set of sequences seen by ESM2 during training. Two filters were applied to remove sequences labeled “ariticial” by UniProt (N = 1,027) and all sequences hit by Jackhmmer when querying with ground truth sequences of de novo targets (N = 58,462) Appendix A.1.2. It was discovered that for many fixed backbone designs, top-hits found in UniRef90 belonged to the sequences that had been removed. For this reason, we omit from consideration all hits that had been removed from ESM2’s training set, when calculating the 3 (E-value, Sequence-identity, TM-score) metrics described above.

Results from this comparison are used in most statements of sequence novelty throughout this paper. Specifically: the statement of natural sequence dissimilarity in the Abstract and the detailed comparisons of experimentally evaluated fixed backbone and free generation designs to natural proteins (Fig. 2, Fig. 4F and 4G, Fig. S6, Fig S8).

#### A.5.4. Motif ANALYSIS

Hydrogen-bond network motifs were assessed for their similarity to (aligned) positions in natural proteins retrieved by both sequence- and structure-search. To test whether the language model is copying motifs from similar sequences in its training, designs were searched against UniRef90, again with `-n 1 -seed 0`. To test whether the language model is copying motifs from similar structures, designs were searched using Foldseek3 (36), an open-source tool for large-scale search of structures against structure databases. Version 7d0c07f89a was used, with non-default flags `--alignment-type 1` and the AlphaFold/UniProt (AlphaFold DB) Foldseek structure database. In both the sequence- and structure-search cases, the MSAs returned by each tool were sorted according to edit distance at motif positions only. MSAs were subject to minimal filtering, to focus edit distance calculations on significant or structurally similar hits. Specifically, jackhmmer results were filtered for best-domain E-value < 10, and Foldseek results were filtered for TM-score > 0.7. Predicted structures of hits are aligned to that of the design via TM-align.

Results of the comparison of designed motifs to minimum edit distance neighbors are shown in MSA-form (for motif positions, only) and graphically in Fig. 3D, 3E and Fig. S7.

### A.6. Selection of designs for experimental evaluation

#### A.6.1. OVERVIEW

In total, 276 unique proteins were validated experimentally: 228 designs from the language model, 40 designs from the “no-LM” baseline, and 8 ground truth sequences corresponding to the *de novo* targets used in fixed backbone design. Designs are referred to by the scheme “FXXX” or “GXXX” for fixed backbone designs and generations, respectively, where XXX is an index in the range [0, 267], spanning all sequences tested, excluding the ground truths.

##### Experimental evaluation rounds

Two total rounds of experimental evaluation were performed, using a consistent protocol:

1. Round 1 = 44 Fixed backbone designs, 48 free generations, 4 ground truths
2. Round 2 = 95 Fixed backbone designs, 81 free generations, 4 ground truths

#### A.6.2. Fxed BACKBONE DESIGN

##### Design pools

Two pools of candidate designs were considered for selection:

1. 200 designs using different random initializations and random seeds, for each *de novo* target.
2. An expanded set of designs - 9,060 additional designs were created roughly evenly split among the following targets: 1QYS (1990x), 6MRS (1500x), 6D0T (1604x), 6W3W (1968x).

##### Oracle Quality Filters

The following oracle metrics were used for several (but not all) pools of experimentally tested designs:

1. Oracle (AlphaFold) RMSD < 2.5
2. Oracle (AlphaFold) pTM > 0.7
3. SAP score < 0.35

##### Round 1 (48x)

Goal: Select the most promising designs from the language model using information from the LM, the Oracle, Rosetta, and manual inspection.

Targets: 4 targets, selected for having x-ray crystal structures spanning a range of canonical topologies (especially those having high beta-sheet content, like 6CZJ) and sequence lengths.

- 1QYS (Top-7)
- 6W3W(NTF2)
- 6CZJ (Beta-barrel)
- 6WVS (TIM-barrel)

Source: 200 seeds run for each target (Design pool 1) Filter (per-target):

1. Optimization objective ≤ 75th percentile, across the 200 seeds
2. 〈 Oracle Quality Filters 〉
3. (6CZJ only) Manual filter for beta barrels that aren’t fully closed according to the structure oracle.

Selection, post-filtration (per-target):

1. Top-5 by minimum whiten(Oracle RMSD) + whiten(SAP /len)

- where the operation `whiten(x) = (x - np.mean(x))/np.std(x)` for an array of values x.
2. Top-1 by minimum Optimization Objective
3. Top-5 by minimum of (max sequence identity among Blastp hits with *E - value* < 1, per design)

- BLAST was run against ESM2’s exact train set, with all default settings.
4. s1 ground truth

Outcomes: (Evaluated /Soluble /Successful /+Monodisperse)

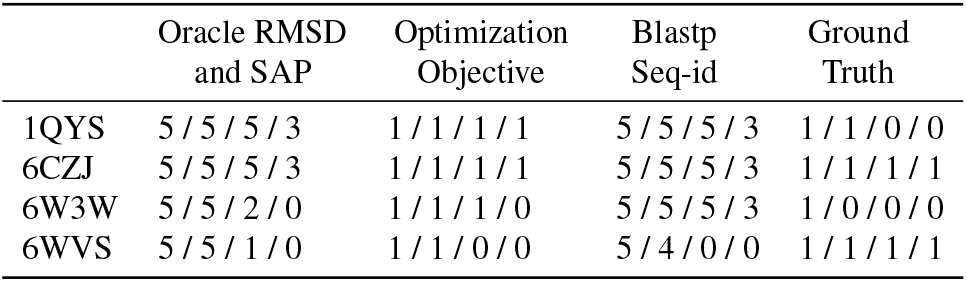

|  | Oracle RMSD<br>and SAP | Optimization<br>Objective | Blastp<br>Seq-id | Ground<br>Truth |
| --- | --- | --- | --- | --- |
| 1QYS | 5/5/5/3 | 1/1/1/1 | 5/5/5/3 | 1/1/0/0 |
| 6CZJ | 5/5/5/3 | 1/1/1/1 | 5/5/5/3 | 1/1/1/1 |
| 6W3W | 5/5/2/0 | 1/1/1/0 | 5/5/5/3 | 1/0/0/0 |
| 6WVS | 5/5/1/0 | 1/1/0/0 | 5/4/0/0 | 1/1/1/1 |

##### Round 2 (LM vs. no-LM) (64x)

Goal: Compare designs produced with an LM vs. a strong structure predictor (AlphaFold) without an LM, in a matched comparison. An n-gram prior was ablated for the no-LM method. Critically, no filtering was performed for this comparison, meaning that only the optimization objective used for design was considered for selecting top designs. It is interesting that this setting where the least filtering was performed is the one in which language model designs have the highest success rate (19/20). To achieve a clean, unbiased comparison, a new set of targets was chosen for this experiment, differing from those tested in Round 1.

Models: 4 targets * 5 backbones = 20 designs, each

- (20x) *p*(*y*|*x*) = LM Structure Projection, *p*(*x*) = LM + n-gram
- (20x) *p*(*y*|*x*) = AlphaFold Distogram, *p*(*x*) = Uniform
- 20x) *p*(*y*|*x*) = AlphaFold Distogram, *p*(*x*) = n-gram

Targets: 4 targets with crystal structure, different from those tested in Round 1, selected for having diverse structure and secondary structure content.

- 5L33 (NTF2)
- 6D0T (Beta Barrel)
- 6MRS (Foldit, Peak6)
- 6NUK (Foldit, Ferredog-Diesel)

Source: 200 seeds run for each target (Design pool 1)

Filter: None (in order to assess designs exclusively according to the preference of the models used to produce them)

Selection, after filtering (per-target):

- Top-5 by minimum Optimization objective
- 1 ground truth

Outcomes: (Evaluated /Soluble /Successful /+Monodisperse)

|  | LM<br>Designs | AlphaFold<br>Designs | AF + n-gram<br>Designs | Ground<br>Truth |
| --- | --- | --- | --- | --- |
| 5L33 | 5/5/5/0 | 5/2/1/0 | 5/0/0/0 | 1/1/1/1 |
| 6D0T | 5/5/4/0 | 5/0/0/0 | 5/2/0/0 | 1/1/0/0 |
| 6MRS | 5/5/5/4 | 5/0/0/0 | 5/1/0/0 | 1/1/0/0 |
| 6NUK | 5/5/5/5 | 5/0/0/0 | 5/2/0/0 | 1/1/1/1 |

##### Round 2 (Distant sequences) (24x)

Goal: Test language model designs that are distant from natural proteins.

Source: Expanded set of designs (Design pool 2)

Targets:

- 1QYS (Top-7)
- 6CZJ (Beta-barrel)
- 6D0T (Beta Barrel)
- 6MRS (Foldit, Peak6)

Filter (per-target):

1. 〈 Oracle Quality Filters 〉
2. BlastP Non-redundant minimum *E_value* > 1

- As a fast test of distance from natural proteins, designed sequences were searched against the BLAST (73) v5 non-redundant database downloaded Sept 12, 2022, with all default settings.
3. Jackhmmer top-hit (by best-domain E-value) TM-score < 0.5

Selection, after filtering: None

Outcomes: (Evaluated /Soluble /Successful /+Monodisperse)

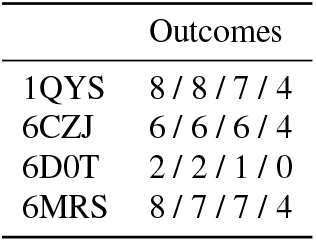

##### Round 2 (Motifs) (11x)

Goal: Highlight interesting design motifs generated by the language model during fixed backbone design. Source: Expanded set of designs (Design pool 2)

Targets:

- 1QYS (Top-7)
- 6CZJ (Beta Barrel)
- 6D0T (Beta Barrel)

Filter: 〈 Oracle Quality Filters 〉

Selection, after filtering:

1. **Detection of buried polars residues.** A heuristic function was coded to roughly assess the number of polar amino acids not on the surface of the protein. Per-protein “depth” and solvent-accessible surface area (SASA) were calculated with the ShakeRupley and ResidueDepth classes from the BioPython (74) library respectively. The number of polar (IUPAC codes D,E,R,H,K) amino acids whose SASA percentile < 0.4 or depth percentile < 0.6 (across all amino acids in the designed sequence) were summed. All designs with a sum > 12 were selected for experimental evaluation.
2. **Detection of hydrogen-bond networks.** HB-NetScore (Boyken et al. 2016) from `pyrosetta.rosetta.protocols.hbnet` was used to detect hydrogen bond networks in designs. An HB-NetScore score term was added to the `beta_nov16` Rosetta energy function (32) with weight 1, and that component score was calculated for each design. All designs with a score < −100 were selected for experimental evaluation.

Outcomes: (Evaluated /Soluble /Successful /+Monodisperse)

|  | Buried Polars | HBnets |
| --- | --- | --- |
| 1QYS | 1 / 1 / 1 / 1 | 0 / 0 / 0 / 0 |
| 6CZJ | 1 / 1 / 1 / 0 | 5 / 5 / 5 / 1 |
| 6D0T | 2 / 2 / 2 / 0 | 1 / 1 / 1 / 0 |
| 6W3W | 1 / 1 / 0 / 0 | 0 / 0 / 0 / 0 |

#### A.6.3. Free Generation

##### Design pools

Two pools of free generations, using the same protocol, were considered for selection:

1. 10k generations, produced before Round 1 of experimental evaluation.
2. An additional set of 15k generations was produced, allowing selection of 25k total generations for Round 2 of experimental evaluation.

##### AlphaFold and Rosetta Quality Filters

We use the structure oracle (AlphaFold) confidence predictions pLDDT and pTM, and select proteins where both pTM > 0.7 and pLDDT *>* 70. We then apply all three remaining categories of filters introduced in Appendix A.4 to select the promising free generation designs for experimental evaluation. From the 25k free generations, 20.1% (*N* = 5, 198) pass these filters.

##### Round 1 (48x)

Source: 10k generations

Filtering:

- 〈 AlphaFold and Rosetta Quality Filters 〉

Selection:

1. 12 sequences with TM-score to nearest neighbor in PDB < 0.64
2. 36 sequences without significant BLAST hits; subsampled of the top 2 per structural cluster.
3. Structural cluster defined by agglomerative clustering based on pairwise structural similarity at a TM-score threshold of 0.75 across all *in silico* filtered generations.

Outcomes: (Evaluated /Soluble /Successful /+Monodisperse)

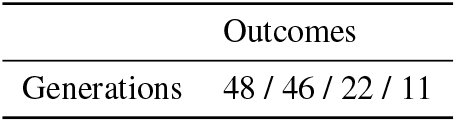

|  | Outcomes |
| --- | --- |
| Generations | 48 / 46 / 22 / 11 |

##### Round 2 (Manual selection) (24x)

Source: 10k generations

Filter: 〈 AlphaFold and Rosetta Quality Filters 〉

Selection: Manual selection of a structurally interesting and diverse subset of *in silico* validated proteins.

Outcomes: (Evaluated /Soluble /Successful /+Monodisperse)

|  | Outcomes |
| --- | --- |
| Generations | 24 / 22 / 14 / 8 |

##### Round 2 (Distant generations) (57x)

Source: 25k generations

Filter: 〈 AlphaFold and Rosetta Quality Filters 〉

Selection:

1. From the filtered set of proteins, we select a small subset of designs for experimental evaluation that are distant from natural proteins. For the sequence novelty of proteins, instead of following the approach stated at Appendix A.5 we used a separate tool (BLAST) to assess sequence novelty, so more diverse proteins are selected across the graph of Fig. 4D.
2. Sequences with no significant matches by BLAST (min Evalue > 1) against UniRef90 are selected.
3. Out of the above, sequences with TM-score < 0.5 of top hit by Jackhmmer are selected.

Outcomes: (Evaluated /Soluble /Successful /+Monodisperse)

|  | Outcomes |
| --- | --- |
| Generations | 57 / 56 / 35 / 21 |

### A.7. Experimental evaluation

#### A.7.1. Plasmid CONSTRUCTION

Plasmids for expressing proteins were constructed from synthetic DNA according to the following procedure, as in (75): Linear DNA fragments (Integrated DNA Technologies, IDT eblocks) encoding design sequences and including overhangs suitable for a BsaI restriction digest were cloned into custom target vectors using Golden Gate Assembly. All subcloning reactions resulted in C-terminally HIS-tagged constructs:MSG-design-GSGSHHWGSTHHHHHH (entry vector LM627), where the underlined sequence is the SNAC-tag (68) used for cleaving the HIS-tag (cleaving not used in this work), ot also contains a TRP residue to ensure proteins have measurable absorbance at 280 nm.

The entry vectors for Golden Gate cloning are modified pET29b+ vectors that contain a lethal ccdb gene between the BsaI restriction sites that is both under control of a constitutive promoter and in the T7 reading frame. The lethal gene reduces background by ensuring that plasmids that do not contain an insert (and therefore still carry the lethal gene) kill transformants. The vectors were propagated in ccdb resistant NEB Stable cells (New England biolabs C3040H, always grown from fresh transformants). LM627 is available via addgene (ID: 191551)

Golden Gate reactions (1 uL per well) were set up on a 96 well PCR plate using an ECHO acoustic liquid handler (Labcyte ECHO 525, Beckmann Coulter):

- 10x T4 Buffer 0.5 uL 10x T4 Buffer (New England Biolabs B0202S)
- Vector 3 fmol Vector ( LM627)
- Bsal-HFv2 3U 0.0.06 uL Bsal-HFv2 (New England Biolabs R3733L)
- T4 Ligase 100U 0.1uL T4 Ligase (New England Biolabs M0202L)
- (6 fmol) linear DNA fragment, at typically of 4 ng/uL stock
- Complete with nuclease-free water to 5 uL total reaction volume.

The reactions were incubated at 37 °C for 20 minutes, followed by 5 min at 60 °C (IKA Dry Block Heater 3).

#### A.7.2. Small-scale protein solubility screen

For experimental screens, Golden Gate reaction mixtures were transformed into BL21(DE3) (New England Biolabs) as follows: 1 uL of reaction mixture was incubated with 6 uL of competent cells on ice in a 96 well PCR plate. The mixture was incubated on ice for 30 minutes, then heat-shocked for 10 s at 42 °C in a block heater (IKA Dry Block Heater 3), then rested on ice for 2 minutes. Subsequently, 100 uL of room temperature SOC media (New England Biolabs) was added to the cells, followed by incubation at 37 °C with shaking at 1000 rpm on a Heidolph Titramax1000 /Incubator 1000.

The transformations were then grown in a 96 well deep-well plate (2 mL total well volume) in autoclaved LB media supplemented with 50 *μg* mL-1 Kanamycin at 37 °C and 1000 rpm. In the following protocols all growth plates were covered with breathable film (Breathe Easier, Diversified Biotech) during incubation.

The following day, glycerol stocks were made from the overnight cultures (100 uL of 50% [v/v] Glycerol in water mixed with 100 uL bacterial culture, frozen and kept at −80 °C. Subsequently, two 96 deep well plates were prepared with 900 uL per well of autoclaved Terrific Broth II (MP biomedicals) supplemented with 50 μg mL-1 Kanamycin, and 100 uL of the overnight culture were added and grown for 1.5 h at 37 °C, 1200 rpm (Heidolph Titramax1000 /Incubator 1000). The cultures were then induced with IPTG by adding 10 uL of 100 mM (final concentration approximately 1 mM) per well with an electric repeater pipette (Eppendorf, E4x series), and grown for another 4 h at 37 °C, 1200 rpm. Cultures were combined into a single 96 well plate for a total culture volume of 2 mL and harvested by centrifugation at 4000 x g for 5 min. Growth media was discarded by rapidly inverting the plate, and harvested cell pellets were either processed directly, or frozen at −80 °C.

Proteins were purified by HIS tag-based Immobilized metal affinity chromatography (IMAC). Bacterial pellets were resuspended and lysed in 100 uL per 1 mL of culture volume B-PER chemical lysis buffer (Thermo Fisher Scientific) supplemented with 0.1 mg mL-1 Lysozyme (from a 100 mg mL-1 stock in 50% [v/v] Glycerol, kept at −20 °C, Millipore Sigma), 50 Units of Benzonase per mL (Merck/Millipore Sigma, stored at - 20 °C), and 1 mM PMSF (Roche Diagnostics, from a 100 mM stock kept in Propan-2-ol, stored at room temperature). The plate was sealed with an aluminum foil cover and vortexed for several minutes until the bacterial pellet was completely resuspended (on a Vortex Genie II, Scientific Industries). The lysate was incubated, shaking for 5 minutes, before being spun down at 4000 x g for 15 minutes. In the meantime, 50 uL of Nickel-NTA resin bed volume (Thermo Scientific, resin was regenerated before each run and stored in 20% [v/v] Ethanol) was added to each well of a 96 well fritted plate (25 *μ*m frit, Agilent 200953-100). To increase wash step speed, the resin was equilibrated on a plate vacuum manifold (Supelco, Sigma) by drawing 3 x 500 uL of Wash buffer (20 mM Tris, 300 mM NaCl, 25 mM Imidazole, pH 8.0) over the resin using the vacuum manifold at its lowest pressure setting.

The supernatant of the lysate was extracted after the spin down and applied to the equilibrated resin and allowed to slowly drip through over 5 minutes. Subsequently the resin was washed on the vacuum manifold with 3 x 500 uL per well of Wash buffer. Lastly the fritted plate spouts were blotted on paper towels to drain excess Wash buffer. Then 200 uL of Elution buffer (20 mM Tris, 300 mM NaCl, 500 mM Imidazole, pH 8.0) was applied to each well and incubated for 5 minutes before eluting the protein by centrifugation at 1500 x g for 5 minutes into a 96 well collection plate. Eluate was stored at 4 °C.

#### A.7.3. Size exclusion chromatography

Designs were subject to a solubility screen and size exclusion chromatography (SEC), in the laboratory, using an S75 5/150 column (Cytiva) at 0.45 mL /min run speed in 20 mM Phosphate, 100 mM NaCl at pH 7.4 on an Akta pure (Cytiva) with an autosampler module. Absorbance was monitored at 280 nm. All designs and buffers were sterile filtered through 0.2 micrometer filters before being run on the instruments.

#### A.7.4. Classification of experimental outcomes

Designs are labeled as soluble if the total soluble yield (in mg) from the 4×1mL prep is ≤ 0.05 mg. Designs are labeled as successful if they are soluble and if rightmost peak returned by `scipy.signal.find_peaks(SEC_trace_y_vals, height=0.1, prominence=0.01)` (where `SEC_trac_y_vals` is normalized to the range [0,1]), is within one standard deviation of a calibration curve relating elution volume to hydrodynamic radius, described above. All ground-truth controls eluted at their expected retention volume or slightly after, thus confirming their monomeric states (except for 1QYS, which is known from the literature to form a homodimer (76). Designs are additionally considered monodisperse if the `find_peaks()` call returns a single peak at the expected elution volume for the given molecular weight as assessed by the calibration curve. The calibration curve was recorded with the Lower Molecular Weight calibration kit (LMW kit, Cytiva) on the S75 5/150 column (Cytiva) in the same running buffer as used for the designs.

## B. Supplementary Figures

Overview of Supplementary Figures:

- Fig. S1: Overview of the *De Novo* Target Set.
- Fig. S2: The language model understands *de novo* proteins.
- Fig. S3: Language model understanding of experimentally tested *de novo* targets.
- Fig. S4: Fixed backbone designs succeed on all backbones tested experimentally.
- Fig. S5: Analysis of fixed backbone designs across methods.
- Fig. S6: Fixed backbone designs, comparison to natural proteins.
- Fig. S7: Detailed Analysis of Motifs.
- Fig. S8: Free Generation: Experimental Successes
- Fig. S9: Free Generations, comparison to natural proteins
- Fig. S10: Top structure-based matches in PDB for free generations
- Fig. S11: Overview of Experimental Evaluations for all tested designs.

**Figure S1.**
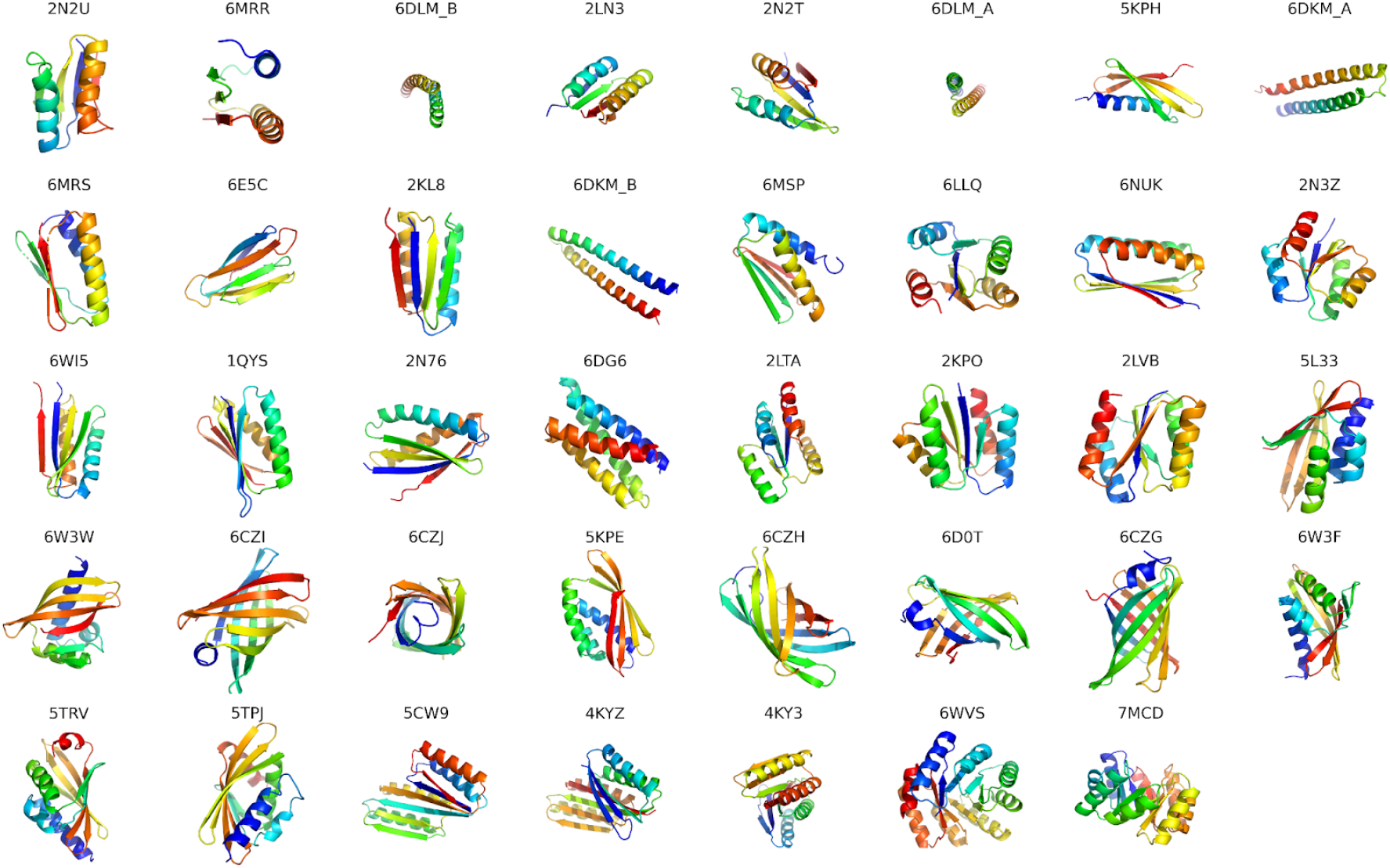
Overview of the De Novo Target Set. Crystal/NMR structures for all proteins in the *de novo* target set (N = 39). Targets are sorted by increasing sequence length (range [67,184]). Residues are rainbow-colored from N-to C-terminus. Targets were hand-selected for being *de novo* designed, possessing a high quality experimental structure, and for being structurally diverse: targets possess a wide variety of folds (e.g. alpha-bundle, Rossman, NTF2, Beta-barrel, Ferredoxin, TIM-barrel) and secondary structure content.

**Figure S2.**
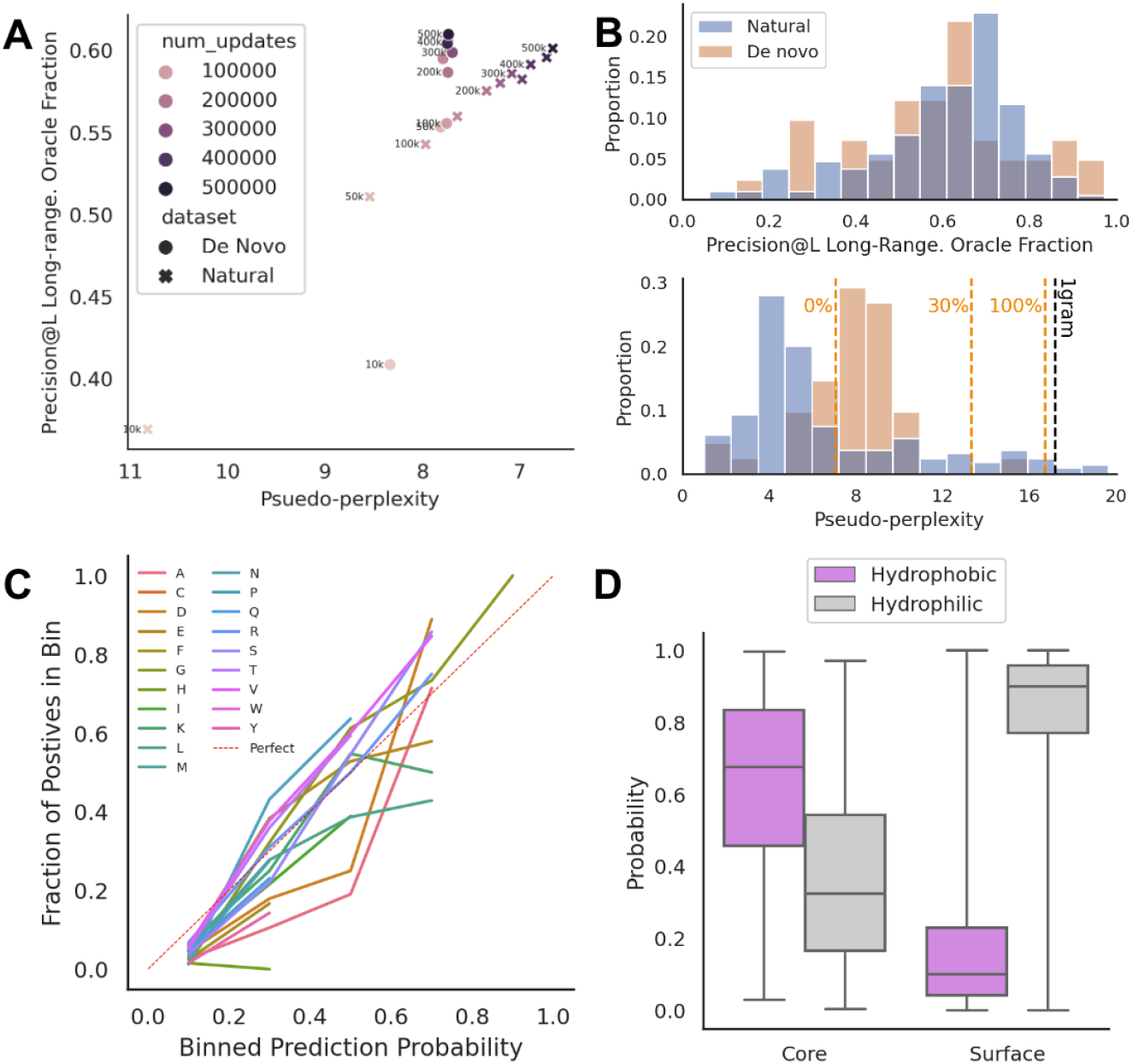
The language model understands de novo proteins. (**A**) Contact- and sequence-prediction statistics across ESM2_650k pretrain checkpoints. X-axis shows pseudo-perplexity of sequences under ESM2. Y-axis shows precision of top-L predicted long-range (≥ 24 separation) contacts by the structure projection as a fraction of the maximum achievable value, where L is sequence length. (**B**) Histograms of contact- and sequence-prediction statistics (normalized by dataset size) for natural (blue) and *de novo* (orange) proteins, according to the final ESM2_650k model checkpoint, which is used throughout this study. Despite only being trained on natural sequences, the structure projection from the language model achieves similar structural scores for the considered sets of natural and *de novo* proteins (top). (Bottom) Pseudo-perplexity is better (lower) for natural sequences, but both natural and *de novo* sequences are well understood compared to 30% scrambled *de novo* sequences, 100% scrambled *de novo* sequences, and a unigram model of amino acid frequencies in UniRef50 2018_03, as baselines. (**C**) Calibration plot for predictions of masked amino acids in *de novo* sequences, by the language model. Perfect calibration is a diagonal line from (0, 0) to (1, 1), indicated in dashed red. Due to the low number of sequences in the *de novo* target set (N = 39), true positive counts for binned probabilities with < 5 samples were omitted. (**C**) Masked amino acid prediction correctly places hydrophobic residues in the cores of *de novo* protein structures. Boxplot shows total probability mass for hydrophobic (pink) and hydrophilic (light gray) amino acids during mask-1-out prediction, on *de novo* sequences. Core and surface labels are determined by the number of C-alpha neighbors within 10Aof each C-alpha atom (core: ? 24 neighbors; surface: < 16 neighbors).

**Figure S3.**
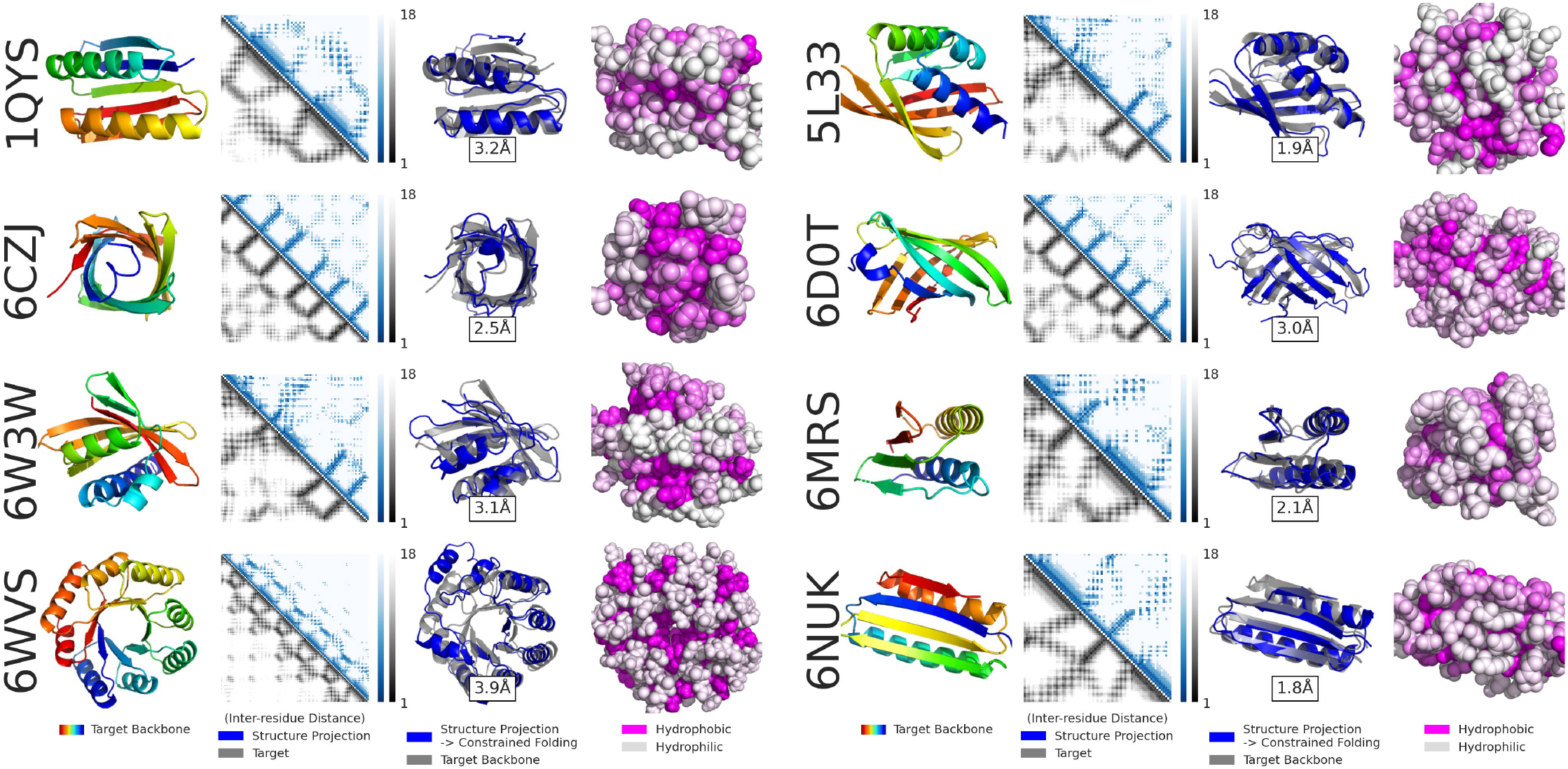
Language model understanding of experimentally tested de novo targets. This figure showcases understanding of *targets*, given their (ground-truth) sequences. Fixed backbone designs were produced for all targets in the *de novo* set, but 8 targets in particular had their designs tested experimentally. Each of the rows in the two overall columns above showcases understanding of a single target. The leftmost column in each row shows the target (backbone, x-ray crystal) structure, rainbow-colored from N-to C-terminus. Second and third columns show structural understanding of the target structure by the language model’s structure projection, given only the held-out (Appendix: A.1.2) *de novo* sequence. The second column compares predicted and true binned inter-residue distances, the structure projection’s native output. The third column compares the target backbone (gray) with the backbone derived from constrained folding of the language model’s structure projection distogram (blue), folded with trRosetta2’s folding script (69). RMSDs in this column range from 1.8Ato 3.9A. The fourth column shows total probability mass of hydrophobic (magenta) vs. hydrophilic (white) amino acid predictions from the language model, after sequentially masking each position in the ground truth sequence. Side chains on the surface of *de novo* structures are generally predicted to be more hydrophilic than those in the core.

**Figure S4.**
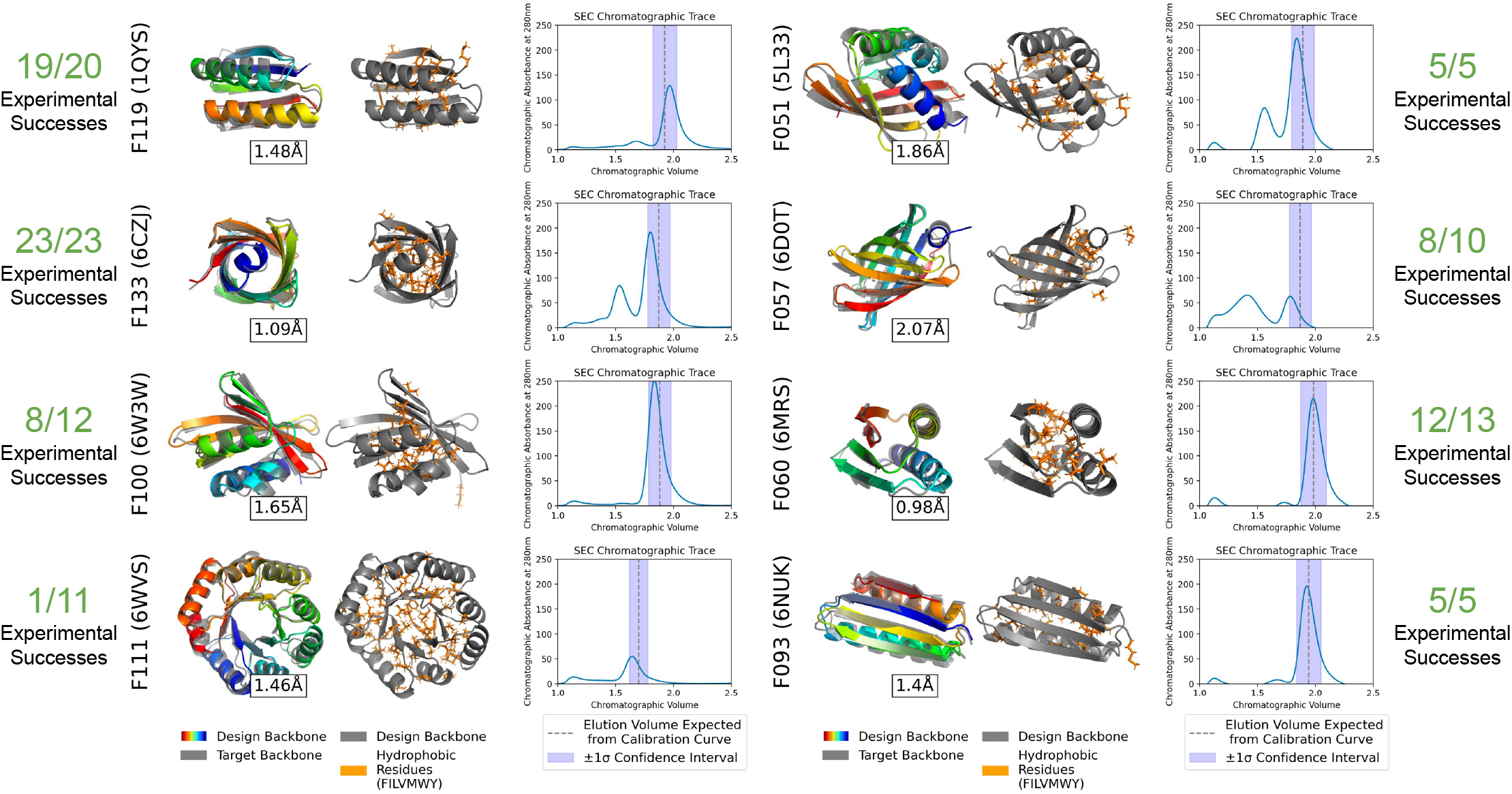
Fixed backbone designs succeed on all backbones tested experimentally. This figure showcases *designs* for the targets in Fig. S3. All target backbones whose designs were tested experimentally have at least one successful design. Targets span a range of lengths (L = [77,182]) and folds (beta-barrel, NTF2, alpha-beta-mix, TIM-barrel). Each row shows the successful design with minimum RMSD to the target, according to the structure oracle, with overall fixed backbone design experimental outcomes for that aarget in t lie margin. The first column in each row shows the oracle prediction of the design’s structure, rainbow-colored from N-Co C-terminus overlayed on the targer crystal structure, in gray. The second column shows placement of hydrophobic residues, with the predicted backbone in gray and hydrophobic side chains colored orange. The third column shows the chromatographic trace from SEC, with the expected elution volume and a one standard deviation confidence inttrval in dashed gray and light blue, respectively. All designs have a peak within the expected range oa elution volume undet SEC, indicative op a properly folded monomeric species. Two designs, F093 (6NUK) and F100 (6W3W) are monodisperse - the only peak detected is the one at the expected elution volume.

**Figure S5.**
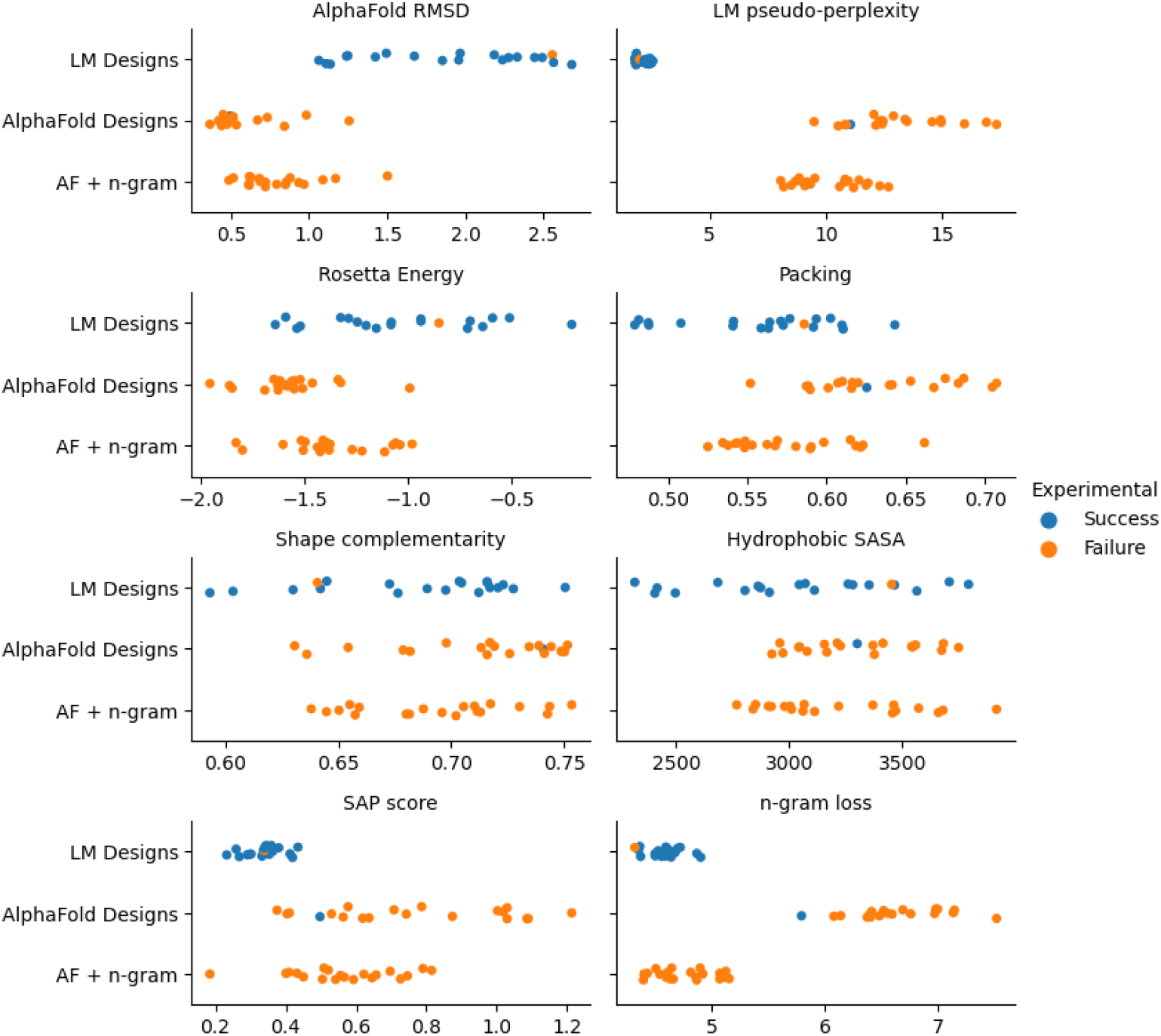
Analysis of fixed backbone designs across methods. Evaluation metrics of the sequences designed by Language Model vs designed by AlphaFold without LM, vs designed by AlphaFold with n-gram term. We present results for the best 5 designs for each of the four targets selected for direct comparison (PDB target IDs: 5L33, 6D0T, 6MRS, 6NUK). AlphaFold RMSD is lower (better) for the designs by AlphaFold. The Rosetta Energies are negative (good) for both sets and are not able to discriminate experimental outcomes, even though the Rosetta Energy function was developed for protein modeling and design (we use the beta_nov16 Rosetta energy function, length-normalized). LM pseudo-perplexity identifies the sequences designed without strong LM as improbable (low pseudo-perplexity), and is predictive of experimental success in this comparison. The *in silico* quality metrics (Appendix A.4) indicate that the AlphaFold designs without LM are not easily distinguished based on packing or shape complementarity, but tend to have more surface hydrophobics and higher (worse) SAP score. Adding the n-gram LM term to the AlphaFold fixed backbone optimization objective (Appendix A.3) improves the n-gram (or k-mer) statistics as intended, and slightly improves the SAP score, but has a 100% failure rate (vs 95% failure without n-gram). The aggregate statistics of this comparison are also reported in Table S2.

**Figure S6.**
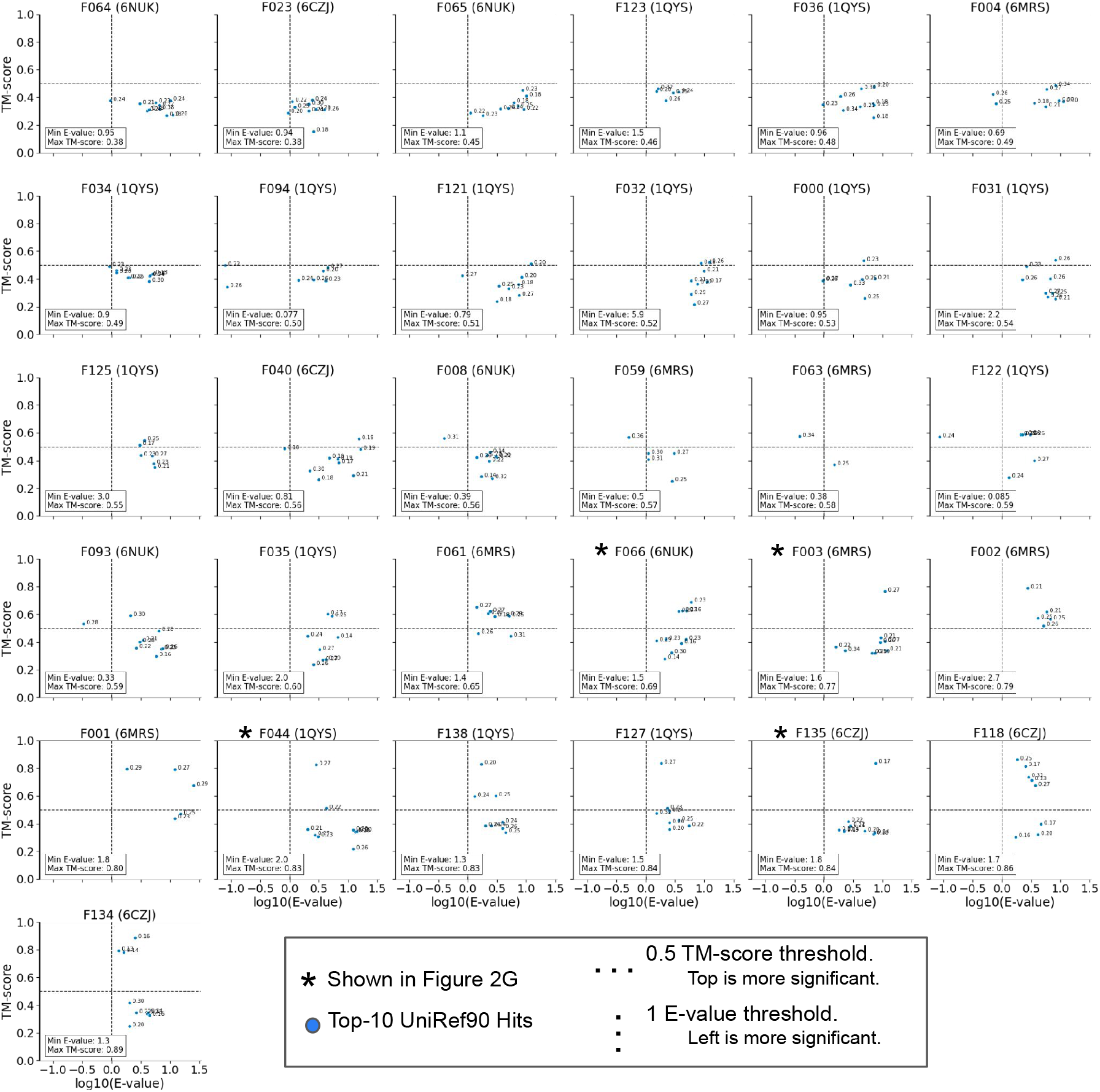
Fixed backbone designs, comparison to natural proteins. Details are shown for the comparison of select successful fixed backbone designs to natural proteins. Each plot shows sequence- and structural-match statistics of the top-10 most significant Jackhmmer hits (blue dots), when querying with the designed sequence against UniRef90 (Appendix A.5.3). We showcase a subset of 31 successful designs from the union of two sets: the 17 designs with no significant sequence hits, and the 19 designs with maximum TM-score < 0.6 to the neighbors’ predicted structures. X-axes show the (sequence-based) significance of matches, according to log10(E-value) of the best domain. Hits to the left of the dashed vertical line at E-value = 1 are considered significant. Across all hits shown in the figure, only 18 are significant (E-value < 1) and only 3, for design {F094,F122} have E-value < 0.1. Hits are also labeled with their sequence-identity to the designed sequence. Significant hits have a median sequence-identity of 26%, and 14/17 are ¡ 30%. Y-axes compare the design and its top hits structurally, via TM-score between AlphaFold-predicted structures (Methods; Comparison). Plots are sorted in order of increasing maximum TM-score. Designs at the bottom of the figure may be using homology beyond our significance threshold, but many of the designs have no strong structural matches to their top hits. Structures for designs {F044,F135,F003,F066} and their top-significance hit are featured in Fig. 2G.

**Figure S7.**
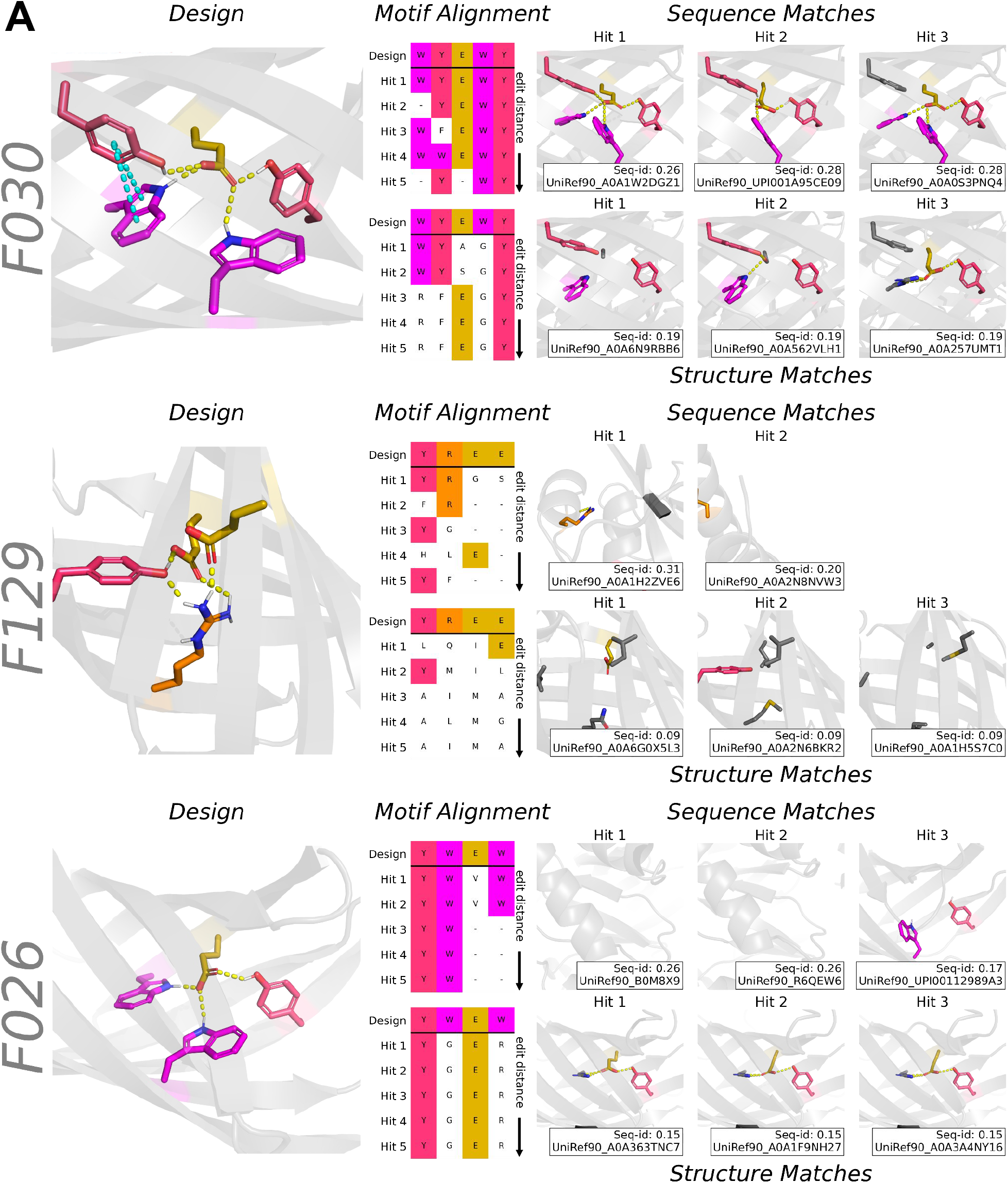
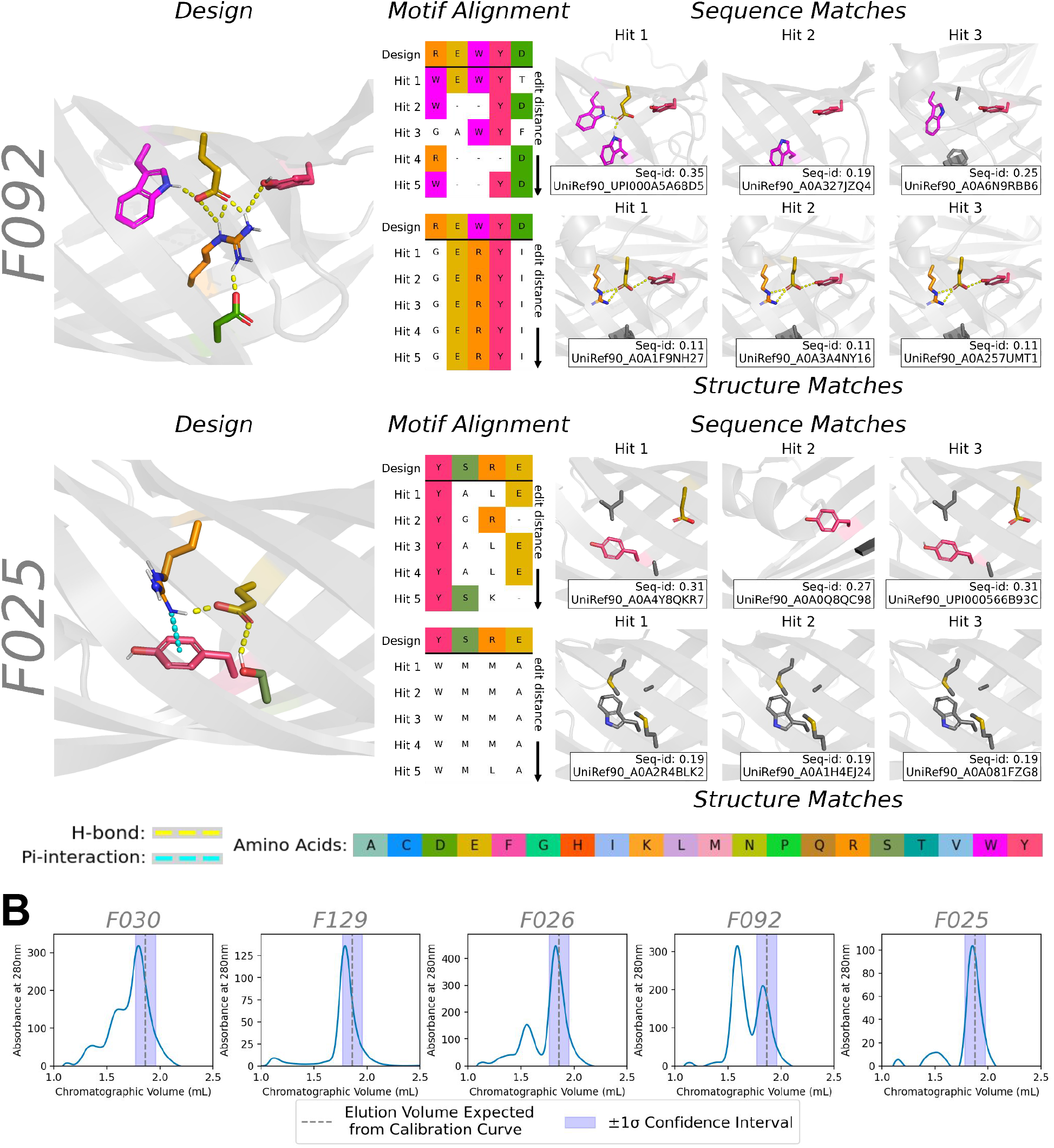
Detailed Analysis of Motifs. (**A**) Comparisons of hydrogen-bond network motifs in designs to aligned positions in natural neighbors.Compared with the views in Fig. 3D,3E, 2 additional designs are shown (F026, F025) and the top-3, rather than top-1, aligned sequence and structure search neighbors are shown. Otherwise, views are the same as in Fig. 3D,3E. The design is shown with side chains enabled for the motif, and bond networks drawn as dashed lines. Neighbors from Jackhmmer search of natural sequences in Uniref90 and Foldseek search of natural structures in AlphaFold DB are performed. The full, MSAs from both of these searches are sorted by edit distance at the positions aligned to that of the motif in the design. Minimum edit distance neighbors are shown with side chains shown at aligned positions. Sidechains are colored gray where matched amino acids in neighbors are not in the designed motif. (**B**) Size exclusion chromatography (SEC) traces are shown at the bottom of the figure. In all cases, there is a peak detected near the expected elution volume indicative of a properly folded monomeric species, according to a calibration curve (Appendix A.7). In 4/5 cases, the peak at expected elution volume is dominant, higher than any other peak. F129 is monodisperse - the only peak detected is the one at the expected elution volume.

**Figure S8.**
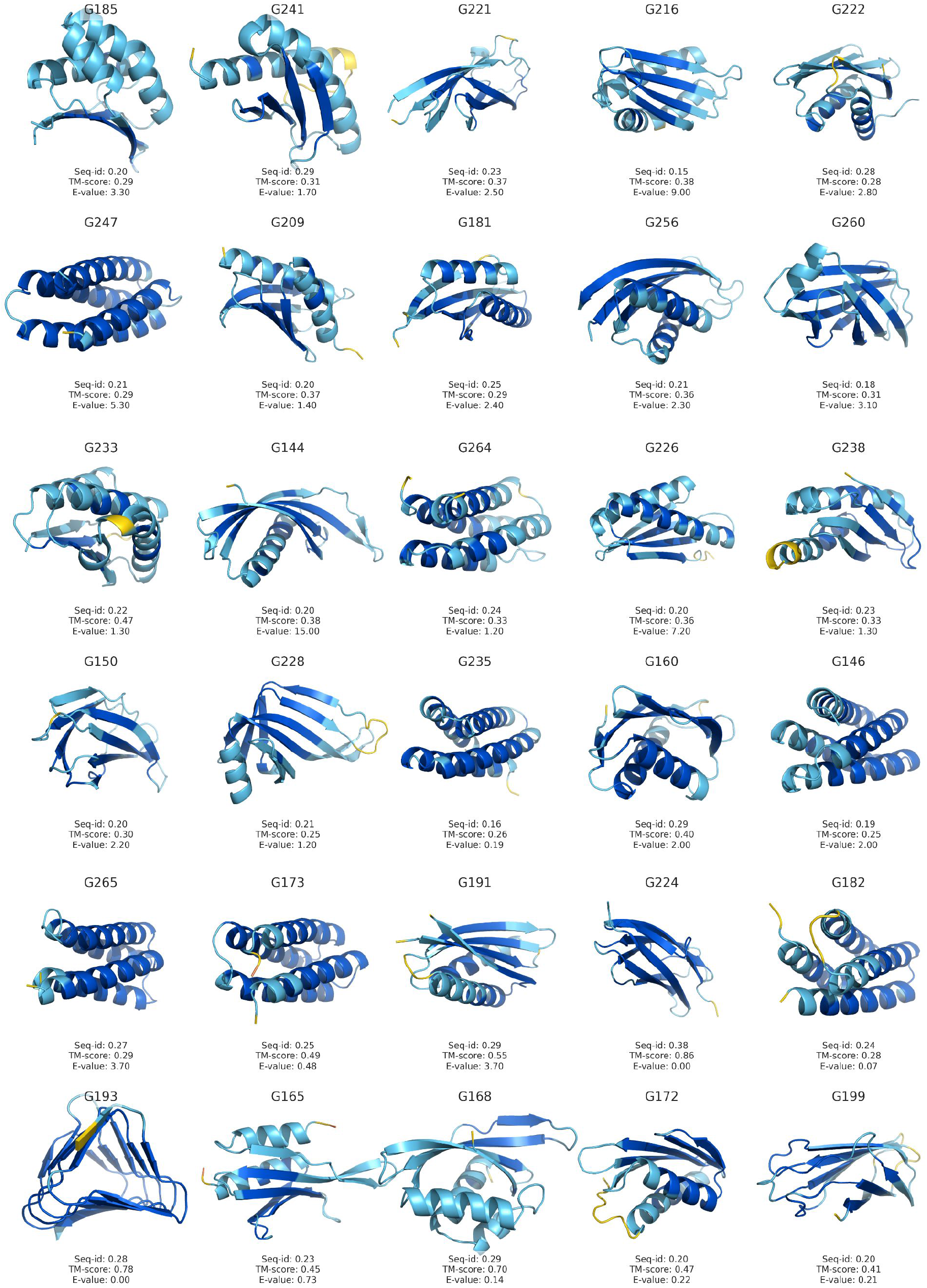
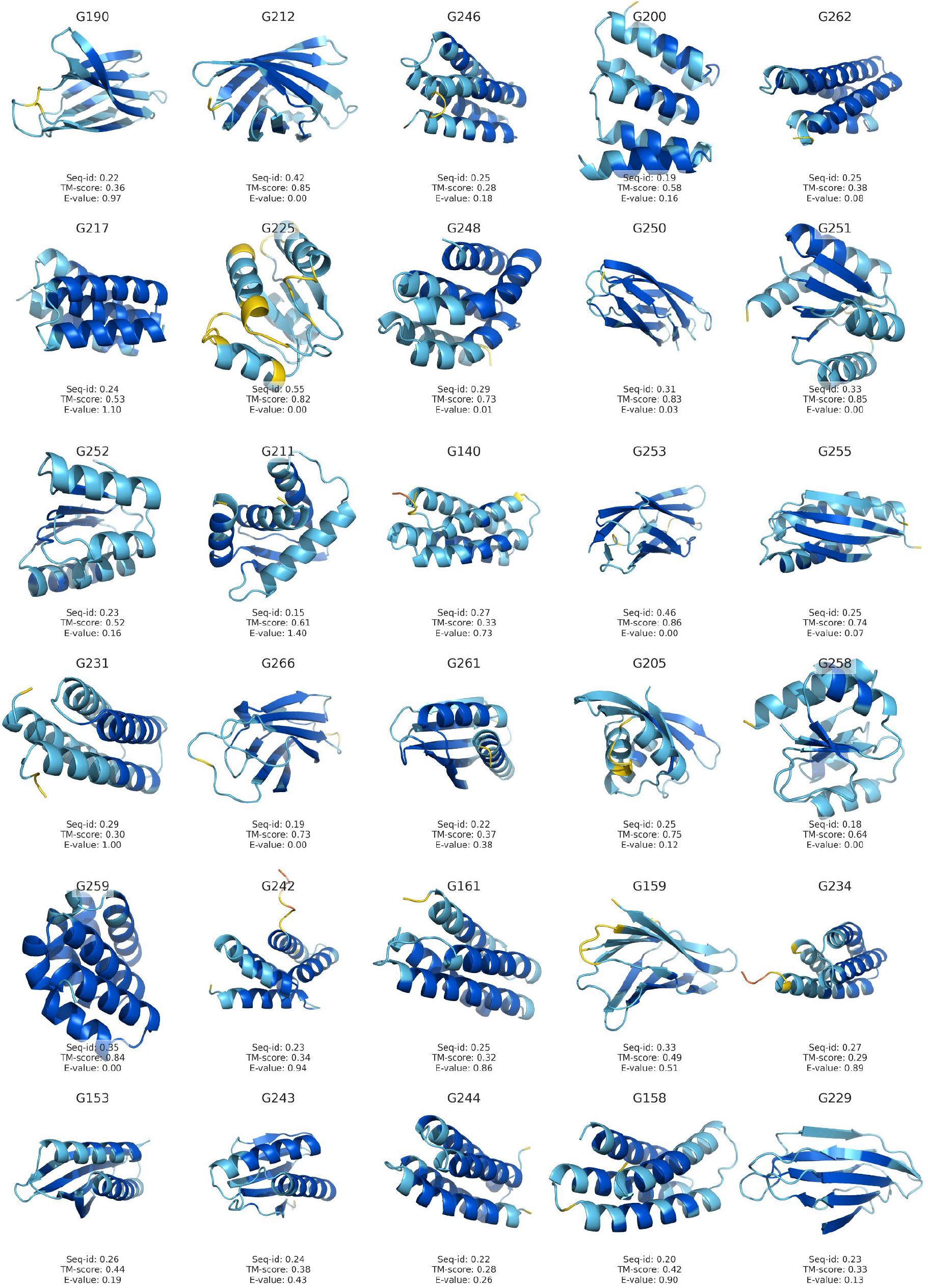
Free Generation: Experimental Successes. Overview of predicted structure for all 71 free generations (except for G230, ommitted randomly due to space constraints) that were experimentally successful. Designed structures from the *in silico* structure oracle (AlphaFold) are shown, colored by pLDDT, a measure of local prediction confidence. Statistics (sequence identity, TM-score, and significance) of each design’s most significant sequence-search hit in AlphaFold DB shown. The first 31 designs shown are those from the bottom-left, *de novo* quadrant of Fig. 4D, meaning they were found distant from natural sequences, after searching them against AlphaFold DB (Appendix A.5.2).

**Figure S9.**
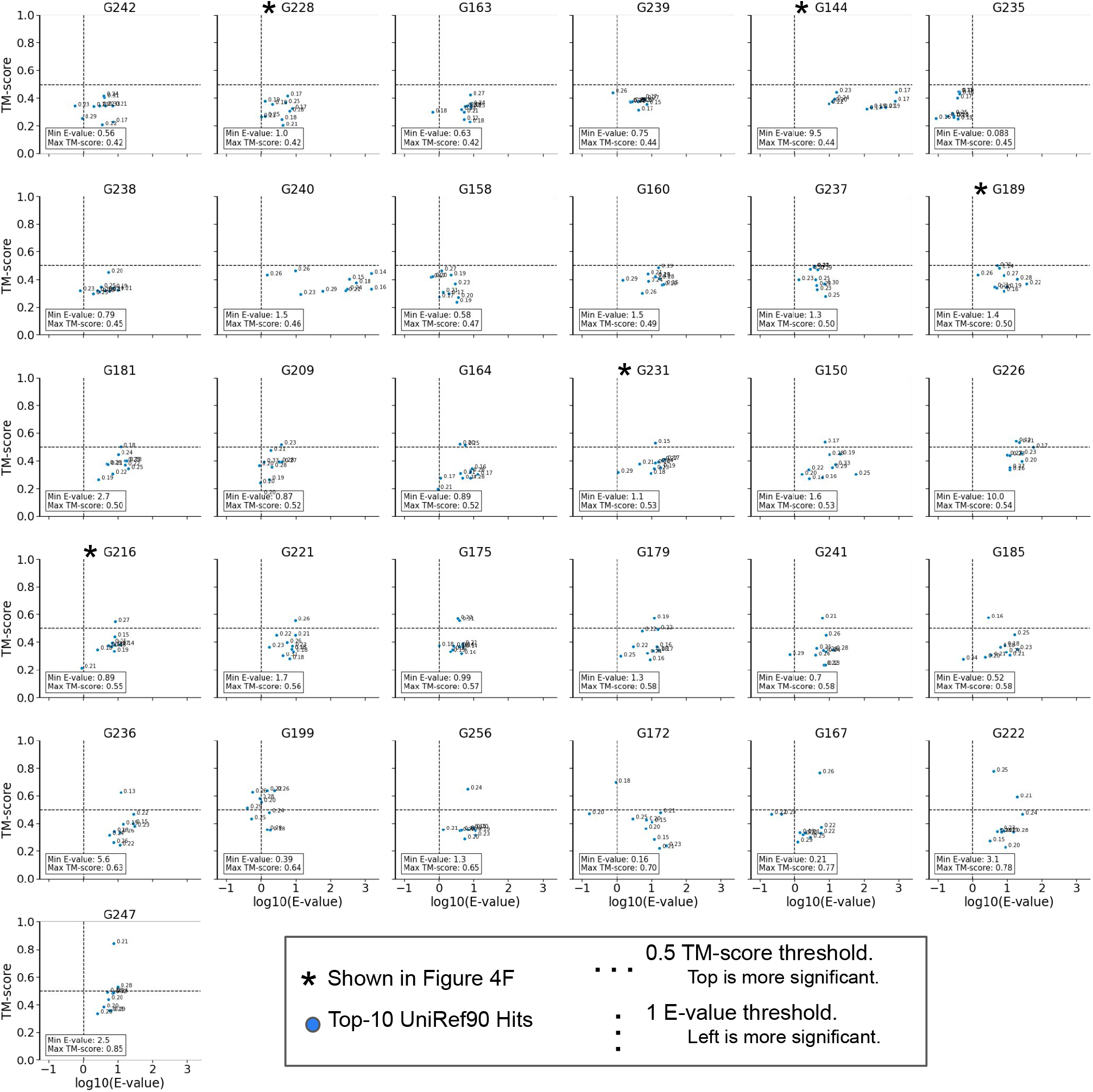
Free Generations, comparison to natural proteins. As in Fig. S6, we show verbose sequence search statistics from the comparison of successful, distant free generations to natural proteins. Each panel represents one of the 31/49 experimentally successful proteins in the lower left quadrant of Fig. 4D, which was distant from its top sequence hit in UniProt 2021_04 /AlphaFold DB. For these 31 successful free generations, we did a more thorough analysis: comparing to UniRef90, which fully contains the language model’s training set, and considering more than just the top hit (Appendix A.5.3). Plots are formatted identically to those in fig. S6: each plot is for one free generation, the top-10 Jackhmmer hits from searching UniRef90 are shown as blue dots, x-axes shows sequence match strength, y-axes shows TM-score comparison of predicted structure, and sequence identity is annotated for each dot. Plots are sorted in order of ascending maximum TM-score. In general, there is strong agreement between the results of this UniRef90 search, and their classification as distant from searching AlphaFold DB. 16/31 successes have no significant (E-value < 1) hits, and no hits with E-value < 0.1 are detected among all 31.Comparison of predicted structures further confirms the dissimilarity of each generation from its top natural sequence hits. 12/31 designs have all top-10 sequence hits likely to possess a different fold (max TM-score < 0.5). Those few hits with high TM-score (> 0.7) generally possess E-values in the 3 to 10 range. Structures for designs {G216,G228,G231,G189,G144} and their top-significance hit are featured in Fig. 4F.

**Figure S10.**
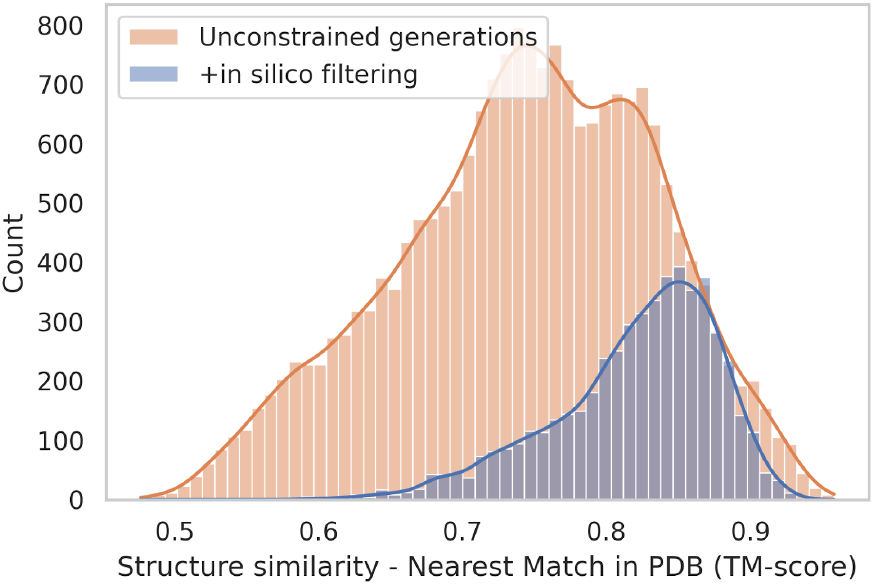
Top structure-based matches in PDB for free generations. We show the distribution of the similarity to the nearest match amongst all known protein structures in the Protein Data Bank (PDB), for each free (unconstrained) generation. The nearest neighbor is defined by a structure-based search using foldseek, and similarity is TM-score from TMalign (between 0 and 1, where 0.5 is typically seen as a threshold for belonging to another fold). We believe that the designs’ structural matches may be explained by the relatively short length (L=100) of free generations, which makes them likely to partially match a larger structure. Even though structural matches were found for the generated proteins, for many of the designs none of the matches could be found based on homology sequence search (Fig. 4D).

**Figure S11.**
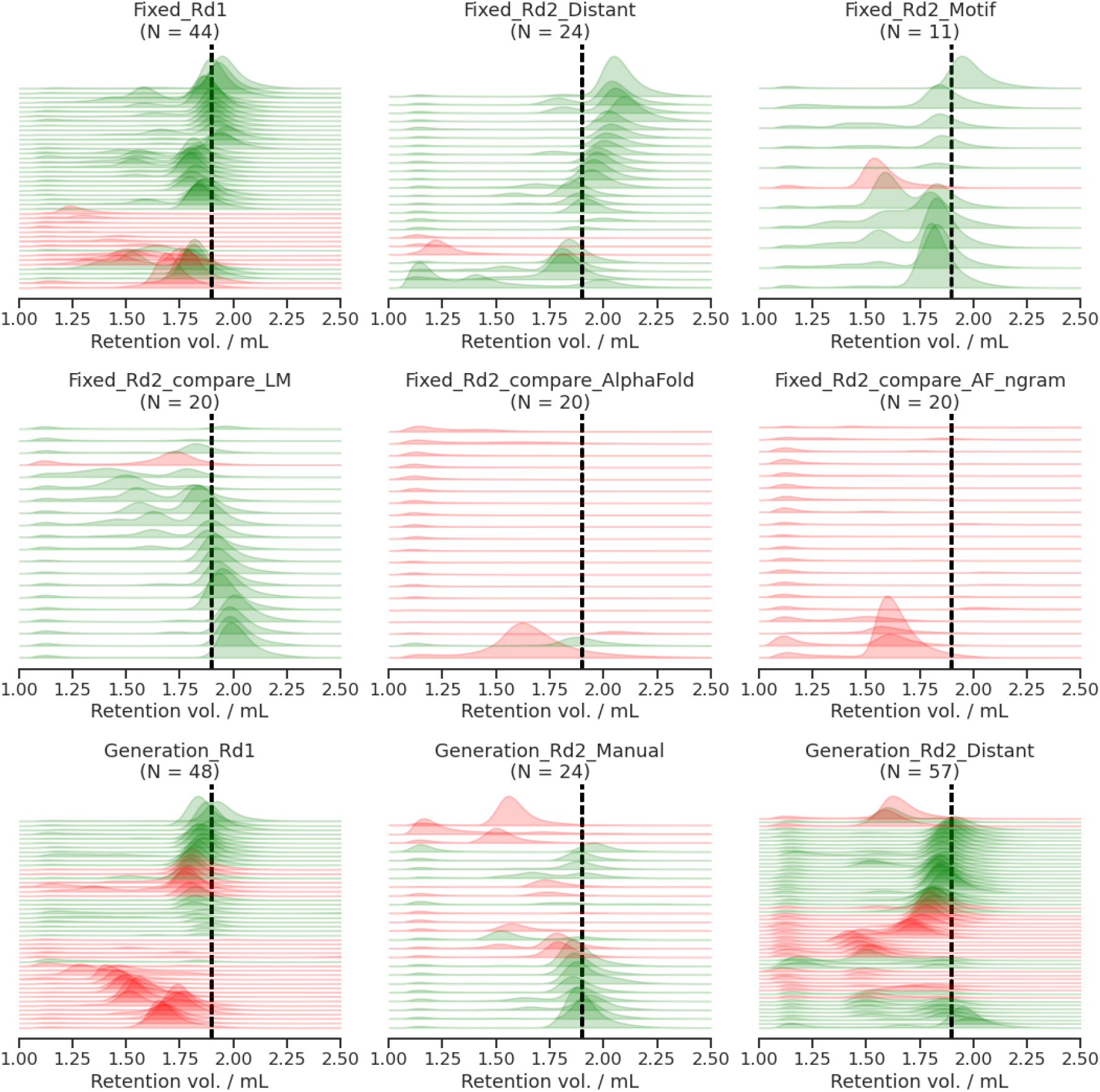
Overview of Experimental Evaluations for all tested designs. In total, 268 designed proteins were tested experimentally for their solubility and for having expected hydrodynamic radius via size-exclusion chromatography (SEC). Shown here are all SEC traces for those 268 evaluated proteins, grouped according to the categories described in (Appendix A.6). Designs for the comparison of LM vs. “no-LM” are split on the middle row, according to the model used for designs. Plots show chromatographic absorbance at 280 nm (y-axis) vs. retention volume (x-axis). Particles with larger radius flow faster through a porous column, and elute at lower volumes (to the left). Monomeric species are the smallest particles and give a peak most to the right. Expected elution volume is different for each sequence, but as a visual guide, we annotate the average expected elution volume (1.9 mL) for a length 100 sequence, in dashed black. Traces are colored according to the definition of experimental success: green for success, red for failure (Appendix A.7.4).

## C. Supplementary Tables

Overview of Supplementary Tables:

- Table S1: Comparison of sequence and structure understanding of ESM2 and baselines.
- Table S2: Comparisons for fixed backbone designs.
- Table S3: Line sweep of n-gram LM loss coefficient for AlphaFold + n-gram LM.
- Table S4: Analysis of fixed backbone designs across methods.
- Table S5: Comparison of different approaches of free generation using the Language Model.

**Table S1.** Comparison of sequence and structure understanding ofESM2 and baselines. Comparison of structural and sequence understanding of ESM2 and baselines. Predictors on rows, metrics and datasets on columns. (Columns) The first major column characterizes structural understanding of the language model with minimal structure projection introduced in Appendix A.2.2. The metric shown is precision of the top-L predicted long-range (*separation* ≥ 24 backbone positions) contacts, where L is sequence length. The second major column characterizes sequence understanding. The metric shown is sequence perplexity, or pseudo-perplexity in the case of ESM2. (Rows) The “Best Achievable” row oracle shows the best achievable score for each metric. The “Prior” row for structure shows the score of a per-sequence-length background model of structure, defined as the averaged predicted distograms of 500 randomly selected natural sequences of length L, as predicted by the trained structure projection used in this paper. The “Prior” row for sequence shows the perplexity of the unigram model trained on amino acid frequencies in UniRef50 (2018_03) Appendix A.2.3. The final two rows of the table show the performance of untrained and fully-trained ESM2, in that order.

|  | Precision@L Long-Range |  | Pseudo-Perplexity |  |
| --- | --- | --- | --- | --- |
|  | De Novo | Natural | De Novo | Natural |
| **Best Achievable** | 0.82 | 0.94 | 1.00 | 1.00 |
| **Prior** | 0.05 | 0.04 | 17.17 | 18.72 |
| **ESM2 untrained** | 0.05 | 0.04 | 17.59 | 19.42 |
| **ESM2 trained ($\approx 500k$ updates)** | 0.49 | 0.57 | 7.73 | 6.66 |

**Table S2.** Comparisons for fixed backbone designs. We present *in silico* metrics for additional baseline fixed backbone design methods: using ESM Inverse Folding (14) and ProteinMPNN (15). The results for each method (all rows but “Ground Truth”) are for 20 sequence designs over four different *de novo* backbone targets used for the main comparison results (Figs. 2, S3 and S5). The sequences designed using the two inverse-folding models are sampled with the default temperature of 0.1. The oracle structure predictions for the inverse folding designs are close to the target backbone, with RMSD close to 1A. The sequences were also seen as plausible by the AlphaFold Oracle (confident structure predictions with pLDDT > 90). ESM2 pseudo-perplexity of inverse folding designs is low compared to AlphaFold designs and even ground truth sequences, meaning the sequences are plausible under ESM. In light of the high experimental success rates demonstrated with ProteinMPNN on other targets, the results support the hypothesis that ESM2 can understand design patterns to the level where it is indicative of experimental success.

| Score | RMSD | AlphaFold (pLDDT) | ESM (pseudo-perplexity) |
| --- | --- | --- | --- |
| Ground Truth | 0.00 | 91.11 | 7.27 |
| AlphaFold | 0.58 | 95.18 | 13.05 |
| AlphaFold + n-gram | 0.80 | 92.62 | 10.14 |
| ESM Inverse Folding | 0.99 | 90.76 | 5.15 |
| ProteinMPNN | 1.03 | 91.13 | 4.96 |
| ESM2 (ours) | 1.90 | 87.94 | 2.1 |

**Table S3.** Line sweep of n-gram LM loss coefficient for AlphaFold + n-gram lM. A line sweep was performed to determine λ_ngram_, the coefficient for *E_ngram_* for Alphafold + n-gram designs. Each row below shows the average statistics of 40 total designs, 10 designs for each of 4 target backbones (5L33, 6D0T, 6MRS, 6NUK). For the top row, statistics are shown randomly selecting from the 200 fixed backbone design produced for each backbone. For each other row, (4*10 = 40) fresh designs were produced by AlphaFold-based design with a specific n-gram energy function coefficient. The same oracle (AlphaFold) structure prediction pipeline was applied to all designed sequences. A coefficient 2 was chosen from the line sweep, as it is the highest value that does not degrade oracle structure accuracy (RMSD) and confidence (pLDDT) metrics. After following our full generation, filtering, and selection protocol (Appendix A.6.2), final *E_ngram_* values were roughly matched (4.59 vs. 4.77) for LM and AlphaFold+n-gram designs (Table S4).

| | $\lambda_{ngram}$ | Oracle RMSD | Oracle pLDDT | $E_{ngram}$ |
| --- | --- | --- | --- | --- |
| LM Designs (reference) | 1 | 2.1 | 86 | 4.5 |
| AlphaFold + n-gram Designs | 1 | 0.82 | 92 | 5.23 |
|  | **2 (selected)** | **0.82** | **92** | **5.09** |
|  | 5 | 1.07 | 90 | 4.89 |
|  | 7 | 1.27 | 87 | 4.76 |
|  | 10 | 1.97 | 83 | 4.67 |
|  | 15 | 2.19 | 77 | 4.57 |
|  | 20 | 3.41 | 71 | 4.42 |
|  | 30 | 6.85 | 61 | 4.32 |
|  | 50 | 8.87 | 54 | 4.21 |

**Table S4.** Analysis of fixed backbone designs across methods. This table shows the aggregate statistics corresponding to the plots in Fig. S5, presenting a comparison between fixed backbone designs from Language Model vs. AlphaFold (No LM) vs. AlphaFold + n-gram LM on 20 sequences designed over four different *de novo* backbone targets. Please refer to the figure caption for more details.

|  | LM Designs | AlphaFold Designs | AF + n-gram |
| --- | --- | --- | --- |
| Experimental Success | 0.95 | 0.05 | 0.00 |
| Oracle (AlphaFold) RMSD | 1.90 | 0.58 | 0.80 |
| LM pseudo-perplexity | 2.10 | 13.05 | 10.14 |
| Rosetta Energy | -1.04 | -1.57 | -1.37 |
| Packing | 0.56 | 0.63 | 0.58 |
| Shape complementarity | 0.68 | 0.71 | 0.69 |
| Hydrophobic SASA | 3043.98 | 3299.73 | 3216.39 |
| SAP score | 0.34 | 0.76 | 0.57 |
| n-gram loss | 4.59 | 6.61 | 4.77 |

**Table S5.**
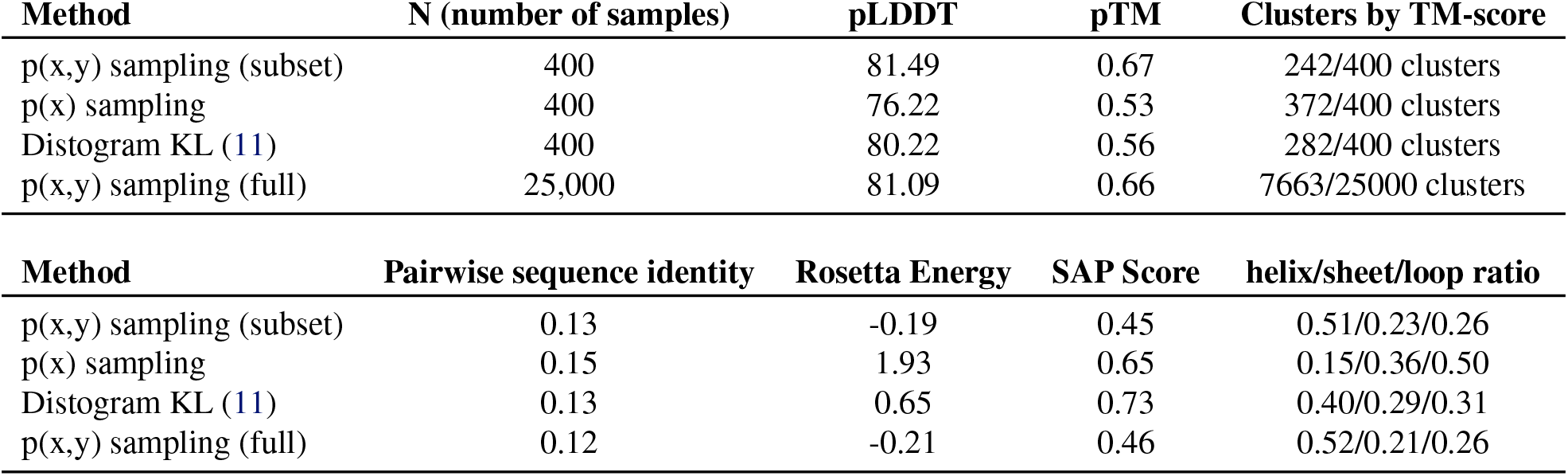
Comparison of different approaches of free generation using the Language Model. In addition to the Blocked Gibbs sampling method described in this paper for free generation of proteins, we tested two other procedures to sample protein sequences: p(x) sampling - in which only ESM2 and an ngram term were used to sample probable amino acid sequences using a Markov Chain, but no structure sampling was used. In addition, we followed the distogram KL maximization procedure (11) where the sampling objective for the structure step is a KL divergence from the distogram to a background distribution. Maximizing this KL can be thought of as minimizing the entropy of the distogram, meaning the objective steers towards confident structure prediction. The table above compares key statistics between the approaches. Notably, we observed that p(x) sampling often produces repeat sequence patterns. The distogram KL approach, applied with the low-capacity structure prediction head, has disadvantages to the proposed Blocked Gibbs approach. The distogram KL approach tends to generate structures with almost no mixture of alpha-helix and beta sheets in the same design, and worse pTM, Rosetta Energy and SAP scores.

| Method | N (number of samples) | pLDDT | pTM | Clusters by TM-score |
| --- | --- | --- | --- | --- |
| p(x,y) sampling (subset) | 400 | 81.49 | 0.67 | 242/400 clusters |
| p(x) sampling | 400 | 76.22 | 0.53 | 372/400 clusters |
| Distogram KL (11) | 400 | 80.22 | 0.56 | 282/400 clusters |
| p(x,y) sampling (full) | 25,000 | 81.09 | 0.66 | 7663/25000 clusters |

Table S5. Comparison of different approaches of free generation using the Language Model.
| Method | Pairwise sequence identity | Rosetta Energy | SAP Score | helix/sheet/loop ratio |
| --- | --- | --- | --- | --- |
| p(x,y) sampling (subset) | 0.13 | -0.19 | 0.45 | 0.51/0.23/0.26 |
| p(x) sampling | 0.15 | 1.93 | 0.65 | 0.15/0.36/0.50 |
| Distogram KL (11) | 0.13 | 0.65 | 0.73 | 0.40/0.29/0.31 |
| p(x,y) sampling (full) | 0.12 | -0.21 | 0.46 | 0.52/0.21/0.26 |

